# Enabling Electric Field Model of Microscopically Realistic Brain

**DOI:** 10.1101/2024.04.04.588004

**Authors:** Zhen Qi, Gregory M. Noetscher, Alton Miles, Konstantin Weise, Thomas R. Knösche, Cameron R. Cadman, Alina R. Potashinsky, Kelu Liu, William A. Wartman, Guillermo Nunez Ponasso, Marom Bikson, Hanbing Lu, Zhi-De Deng, Aapo R. Nummenmaa, Sergey N. Makaroff

## Abstract

Modeling brain stimulation at the microscopic scale may reveal new paradigms for various stimulation modalities. We present the largest map to date of extracellular electric field distributions within a layer L2/L3 mouse primary visual cortex brain sample. This was enabled by the automated analysis of serial section electron microscopy images with improved handling of image defects, covering a volume of 250 × 140 × 90 μm^3^. The map was obtained by applying a uniform brain stimulation electric field at three different polarizations and accurately computing microscopic field perturbations using the boundary element fast multipole method.

We used the map to identify the effect of microscopic field perturbations on the activation thresholds of individual neurons. Previous relevant studies modeled a macroscopically homogeneous cortical volume. Our result shows that the microscopic field perturbations – an ‘electric field spatial noise’ with a mean value of zero – only modestly influence the macroscopically predicted stimulation field strengths necessary for neuronal activation. The thresholds do not change by more than 10% on average. Under the stated limitations and assumptions of our method, this result justifies the conventional theory of “invisible” neurons embedded in a macroscopic brain model for transcranial magnetic and transcranial electrical stimulation.

However, our result is solely sample-specific and largely neglects the effect of the microcapillary network. Furthermore, we only considered the uniform impressed field and a single-pulse stimulation time course.

**Significance statement:** This study is arguably the first attempt to model brain stimulation at the microscopic scale, enabled by automated analysis of modern scanning electron microscopy images of the brain. It concentrates on modeling microscopic perturbations of the extracellular electric field caused by the physical cell structure and is applicable to any type of brain stimulation.

**Data availability statement:** Post-processed cell CAD models (383, stl format), microcapillary CAD models (34, stl format), post-processed neuron morphologies (267, swc format), extracellular electric field and potential distributions at different polarizations (267x3, MATLAB format), *.ses projects files for biophysical modeling with Neuron software (267x2, Neuron format), and computed neuron activating thresholds at different conditions (267x8, Excel tables, without the sample polarization correction from Section 2.8) are made available online through **BossDB**, a volumetric open-source database for 3D and 4D neuroscience data.

## 1. Introduction

Advances in brain stimulation leverage models of electric fields within the brain for understanding the brain’s response to external stimuli provided by transcranial magnetic stimulation (TMS) [1], transcranial electrical stimulation (TES) [2], deep brain stimulation (DBS) [3], electroconvulsive therapy (ECT) [4], or similar excitation. However, accurately modeling these fields within the brain presents a great challenge due to the brain’s complex microstructure, including the intricate networks of neurons and blood vessels [5],[6],[7],[8],[9] all of which must be included into the model.

The conventional approach of field modeling in brain stimulation [10],[11],[12],[13],[14],[15],[16] ignores all microscopic structures, with each region represented by macroscopically averaged properties (e.g., isotropic or anisotropic macroscopic conductivity). Only after the macroscopic electric field is evaluated with finite element method (FEM) [17], are isolated neurons “reinserted” to model their biophysical response.

Ignoring microstructure is a recognized limitation but treated as a *fait accompli* because of known computational intractability of FEM for very complex geometries, except for models of isolated and simplified axonal and vascular geometries (cylinders, spheres, regular polygons, etc.), where the importance is demonstrated both theoretically and experimentally [18],[19],[20],[21],[22],[23],[24],[25],[26].

Over the last few years, several brain samples with realistic anatomical microstructure enabled by automated analysis of serial section electron microscopy images were made available. Those are an L2/L3 mouse primary visual cortex brain sample (250 ×140×90 μm^3^ volume) or IARPA Phase I [5],[6],[7] with both membrane meshes and morphological reconstructions, a 1 mm^3 mouse primary visual cortex brain sample with membrane meshes and cortical column reconstruction or IARPA Phase III [7],[8], and a 1 mm^3 human brain sample (temporal cortex) with membrane meshes and some morphological reconstructions [9]. All samples are augmented with capillary blood networks (about 230 millimeters of blood vessels in [9]).

Is it possible to electromagnetically model these samples to extract information useful for brain stimulation? The present study introduces arguably a first attempt to do so. In contrast to the common FEM, we utilize a matrix-free iterative boundary element fast multipole method (BEM-FMM) [27],[28],[29] with a new nested iterative solution based on physical separability of different cells. The BEM-FMM charge-based formulation [30],[31],[32],[33] could presently handle up to ∼200 million surface degrees of freedom simultaneously. It is readily applicable to computing electric charges induced on nonconducting cell membranes (under initial polarization) of realistic closely spaced neuronal cells and nonconducting (to the main order of approximation) walls of microcapillaries. These charges generate their own secondary electric field ***E***^***s***^ which alters the primary macroscopic brain stimulation field ***E***^***i***^ (due to a TMS coil for example) and thus changes expected activating threshold of the neurons.

In this approximation, the extracellular solution becomes fully decoupled from the intracellular solution. For a finite brain sample subject to an impressed field ***E***^***i***^, the extracellular solution is a unique solution of the exterior Neemann problem [37],[38]. Field variations on the cellular level result in an ‘electric field spatial noise’ with a mean value of zero. In contrast to temporal noise studies (cf. [39],[40]), such a noise is large and rather multiplicative since microscopic field perturbations are directly proportional to the local values of the impressed field ***E***^***i***^ itself.

As a result, a map of distributions of the extracellular electric field within the entire brain sample has been created. The electric field computations alone required approximately 3 months of continuous CPU time, utilizing 7 modern terminal servers simultaneously as well as multiple nodes of a high-performance computing cluster. We used the map to identify microscopic perturbations of the extracellular electric field and their effect on the activating thresholds of individual neurons. Previous relevant studies modeled a macroscopically homogeneous cortical volume.

To do so, we averaged the computed total extracellular field over membrane cross-sections and obtained a microscopically altered 1D activating function [41],[42],[43] at every cross-section of a neuronal process, in the approximation of the 1D cable equation. Thus, we did not model the volumetric intracellular space and did not employ the true bidomain approach [21],[44]. This is because the length-to-diameter ratio for an axonal arbor (most sensitive to activation) of a realistic cortical neuron maybe as high as 60,000 (based on data from [7],[8][9]), which facilitates the use of the 1D cable model.

The extracellular electric field noise defined above alters the activating function, potentially causing local membrane depolarization and inward transmembrane current injection at its positive peaks, while nothing flows at the hyperpolarization peaks – the noise is being rectified. Therefore, our initial hypothesis was that microscopic extracellular noise substantially eases neuron activation as compared to microscopically homogeneous brain modeling.

The question we would like to answer here is: by how much? To do so, standard biophysical analysis with Neuron software [45] was further carried out based on neuron morphologies precisely linked to the physical neurons and known ion channel models for rodents [46],[10],[11],[12]. The brain sample under study was considered as it stands i.e., no cloning, elongation, or scaling neuronal processes/capillaries of any kind are made.

The study is organized as follows. Section 2 Materials and Methods describes assembly of the computational brain sample model (further detailed in Supplements A, B), describes computational workflow for the extracellular field (further verified in Supplement E) as well as the biophysical modeling setup (further detailed in Supplement C). Section 3 Results presents major computational findings for corrected activating thresholds with other results reported in Supplement C. Section 4 Discussion summarizes and explains practical findings of the study and states limitations of our approach. Section 5 Conclusions concludes the paper.

## 2. Materials and Methods

### 2.1 Brain sample – MICrONS L2/L3 dataset of mouse visual cortex

The brain sample is a semi-automated reconstruction of the L2/3 P36 male mouse primary visual cortex volume of approximately 250×140×90 μm in size (Fig. 1a-d) from electron microscopic images with a resolution of 3.6×3.6×40 nm [5],[6],[7]. This reconstruction includes physical (not skeletonized) pyramidal and non-pyramidal neurons, astrocytes, microglia, oligodendrocytes and precursors, pericytes, vasculature, nuclei, mitochondria, and synapses [5],[7]. In total, 364 excitatory (Fig. 1e) and 32 inhibitory (Fig. 1f) neurons with cell bodies within the volume (Fig. 1c) were segmented, along with some microvasculature (Fig. 1d) [7]. For 301 neurons, the labeled neuronal centerlines (skeletons) in the common `swc` format were also made available [7]. This provides a unique opportunity to estimate the effect of the local field perturbations using biophysical analysis [45].

**Fig. 1.**
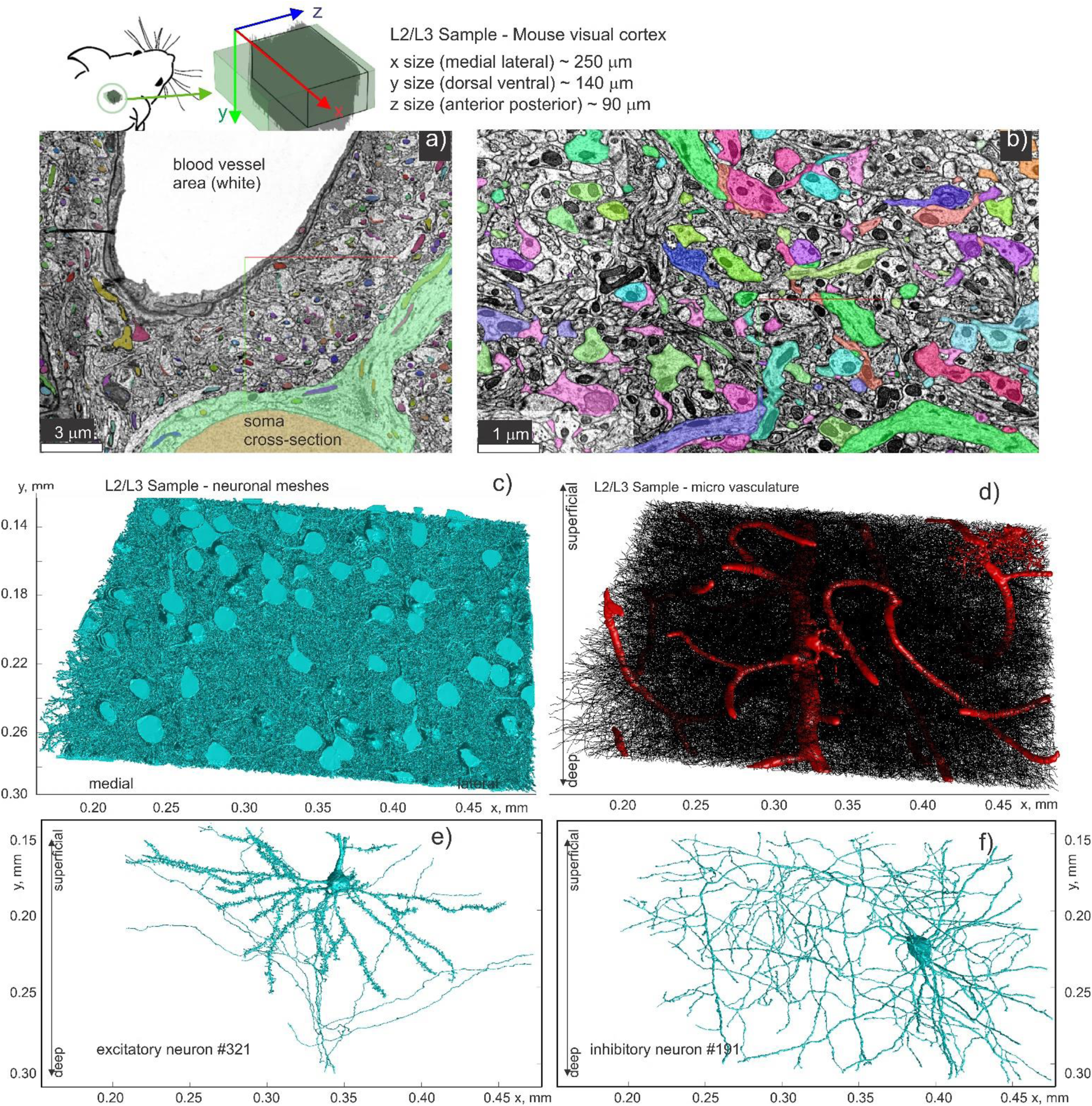
Structure of L2L3 sample under study. a) Snapshot of relative position of a neuron (soma) vs. a nearby blood capillary (white area) within the sample [7], [5]. b) Snapshot of different neuronal processes labeled with different colors at a higher resolution. c) Segmented neuronal bodies of the entire sample. d) Centerlines/skeletons of individual neurons and segmented blood vessels in red (with an astrocyte at top right). The blood network of this sample was not the major target and is severely underdeveloped. e,f) Excitatory (pyramidal) and inhibitory (basket) neurons within the sample volume to scale. The axes differ from the mouse-centric coordinate system [5].

### 2.2 Construction of computational cell meshes. Volumetric fraction of neurons

Only the neurons themselves and the available vasculature (with a few astrocytes) – the largest perturbers of the impressed or primary electric field ***E***^***i***^ – were included into the present numerical analysis. Numerous neuroglia cells seen in Fig. 1a,b were not included in the analysis due to their small size, which made them impossible to model, even with advanced numerical simulation tools. The volume occupied by these microstructures (extracellular space) was considered approximately homogeneous. The volumetric fraction of the neuronal cells within the brain volume was estimated in Table S1 of Suppl. A as 30±5% by computing the volume of each neuron and taking into account neuronal mesh intersections (below). This is significantly less than the previously reported values on the order of 70% [48],[49],[50] (Suppl. A). One reason for it is that all neurons with somas outside of this relatively small sample were omitted and their dendritic/axonal processes, which are still within the sample, were ignored during the segmentation.

Supplement A describes the post-processing procedure for the original sample and its results in detail. Based on this data, Table S1 of Supplement A lists major topological and biophysical parameter of the created computational testbed, including estimated sample anisotropy. Both the facet size of the triangular membrane discretization (Fig. 2a) for the electromagnetic analysis as well as the skeleton’s spatial resolution (Fig. 2b,c) for the biophysical analysis were kept at approximately 100 nm to resolve large yet very localized variations of the impressed field due to multiple nearfield interactions of the neuronal processes as well as multiple intersections of neuronal arbors of different neurons (below).

**Fig. 2.**
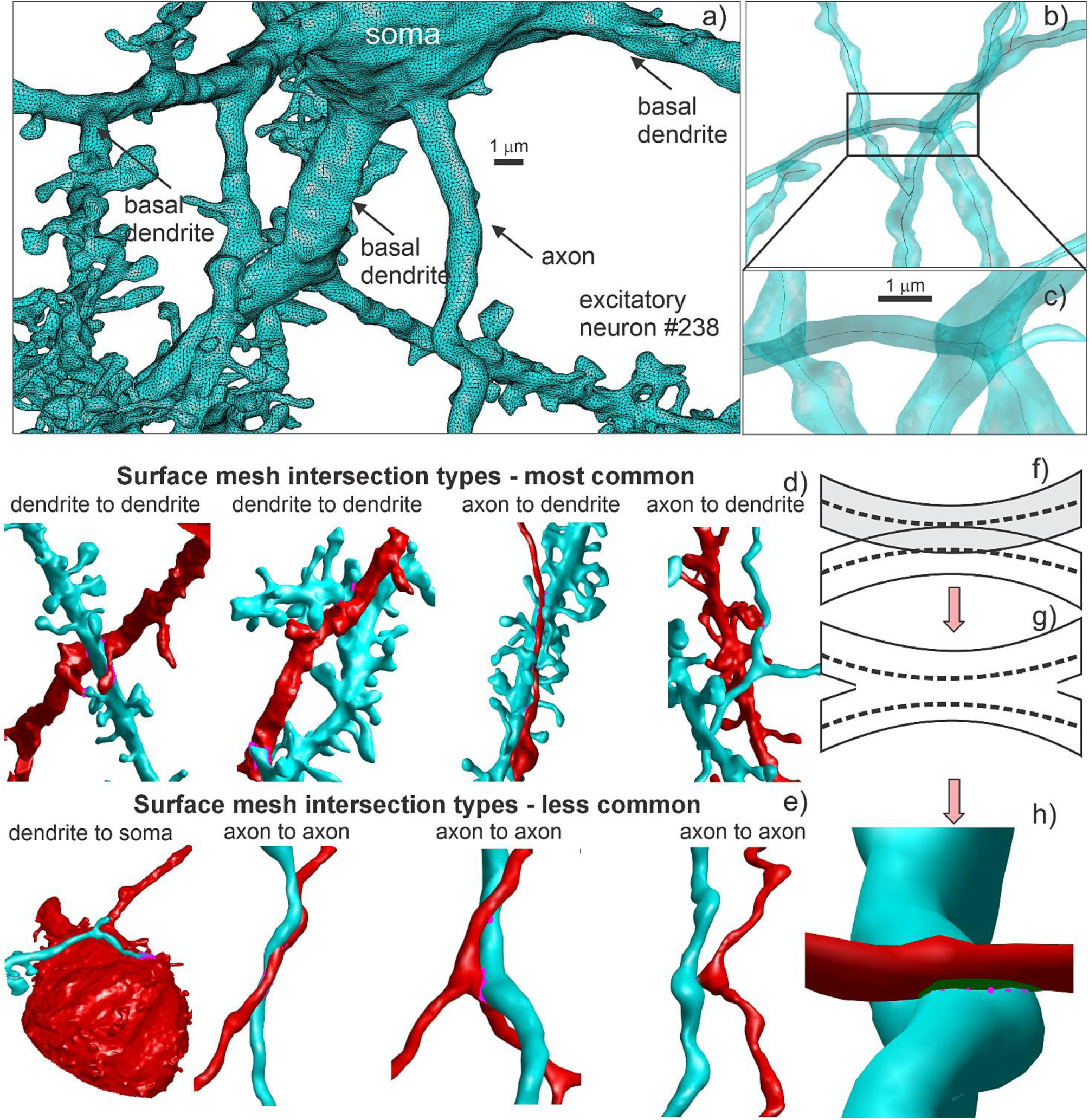
a) Final high-quality membrane surface triangular mesh for one neuron (#238) which is located approximately in the middle of the sample. The membrane mesh has 2 million facets in total and a 100 nm resolution. b,c) Healed centerlines of a neuron. d) Most common intersections observed – dendrite to dendrite and axon to dendrite intersections. Two different colors correspond to the two different neurons. e) Less common intersections – dendrite to soma and axon to axon, respectively. f,g) Mesh intersection resolution via removal of inner triangles. h) Result of this operation for an axon-to-dendrite intersection. Removed triangular facets are marked dark green.

### 2.3 Neuronal mesh intersections (synaptic interconnections) and their resolution

The neuronal and vasculature surface meshes so constructed were found to heavily intersect with each other at multiple locations (cf. Fig. S2 of Suppl. A). Some of these intersections reflect true synaptic interconnections between different neurons with a list of approximately 2,000 synaptic clefts quantified in [6] for the same sample. At the same time, many intersections are likely due to the imprecise segmentation and subsequent, not entirely identical, remeshing. Fig. 2d illustrates the most common intersections – interconnections – including the dendrite to dendrite and axon to dendrite intersections. Two different colors correspond to the two different neurons. Fig. 2e illustrates the less common intersections – dendrite to soma and axon to axon, respectively. Additionally, Supplement A demonstrates an intersection map between neighboring excitatory neurons #238 and #290, respectively (Fig. S2). Fig. S2 also presents a mesh intersection roster of excitatory neuron #238 with all other neurons of the sample. It is observed that neuron #238 intersects with almost all neurons of the sample. Similar results were observed for other larger neurons within the sample.

To resolve the mesh intersections numerically, we used the method illustrated in Fig. 2f,g,h. For two intersecting processes from Fig. 2f, all facets belonging to either process and having at least one node within another process (its surface mesh) have been removed. This operation does not exactly join the two meshes but leaves a narrow gap (one the order of triangle size or ∼100 nm) in between as seen in Fig. 2h. When the gap size is small compared to the intersection domain itself, which is achieved by a present fine mesh resolution, the gap has a minor influence on the results as demonstrated by our refined mesh computations (Supplement A). Note that this method removes coincident or interpenetrating membranes and joins the corresponding intracellular spaces.

### 2.4 Computing extracellular electric field and extracellular electric potential

Our electromagnetic modeling setup only computes the extracellular electric field in a finite brain specimen. For the computation of the extracellular electric field, the approach developed in two relevant yet smaller-scale studies [32],[33] has been used. It assumes that the membrane is non-conducting (insulating) at the end of initial polarization [34],[55],[56], during the application of a relatively short (0.1-1 ms or so) brain stimulation pulse. The same condition applies to the walls of the blood capillaries. The condition of the insulating membrane makes it possible to fully decouple the extracellular solution in the form of a classic uniquely solved exterior problem with the simple Neumann (zero normal current density) boundary conditions at the outer surface of the membrane. In response to an impressed or primary electric field ***E***^***i***^ with the (local) potential *φ*^***i***^, induced surface charges will accumulate on the membrane surfaces. The secondary field of these charges, ***E***^***s***^, will alter the extracellular electric field. The total field in the extracellular space will become ***E***_e_ = ***E***^***i***^ + ***E***^***s***^ and the total extracellular potential will become *φ*_e_ = *φ*^***i***^ + *φ*^***s***^. The goal of the electromagnetic part of this modeling pipeline is to find the secondary field and potential. We have applied a uniform electric field ***E***^***i***^ with the amplitude of 100 V/m to the entire sample for each independent field polarization (*x, y, z*).

We assumed that the sample, which has the form of the non-orthogonal parallelepiped (Fig. 1c), is embedded into a homogeneous medium with the same extracellular conductivity. In this formulation, the exact conductivity value does not matter, and no box around the irregular sample is needed.

However, this formulation still needs adjustment to match the impressed uniform electric field outside the sample with minimal global distortion. In other words, the finite microscopic sample must be embedded into a conventional microscopically homogeneous brain model with minimal distortion. There are different ways to accomplish this. One way is to increase extracellular conductivity within the finite specimen. This will lead to lowering the average extracellular electric field within the specimen. The method for determining this adjustment is based on modeling an artificial rectangular-cuboid sample that mimics the original specimen. This method is described in detail further in this section.

### 2.5 Application of BEM-FMM. Nested iterative algorithm

The single-layer charge formulation for the exterior Neumann problem allows for direct application of the charge-based boundary element fast multipole method (BEM-FMM) developed previously [30],[31],[32],[33]. The BEM-FMM is ideal for this type of problem since it does not require construction of an explicit system matrix and does not require a volumetric mesh for the extremely complicated interior/exterior domains. For the present case, however, the iterative BEM-FMM was only capable of computing the field of ∼100 interacting neurons (with ca 0.2 billion facets) simultaneously. For larger samples, it required a prohibitively large RAM more than 1 TB and was experiencing slower convergence of the underlying GMRES algorithm [60].

Therefore, a nested iterative method has been developed and is illustrated in Fig. 3a. At initial step 1, each full neuron is solved in isolation and is not affected by charges deposited on other neurons. The membrane charge deposition in response to the impressed field ***E***^***s***^ (or the self-interaction) is found via the standard iterative BEM-FMM solution for every neuron in homogeneous space, with other neurons still no existing. Local charge density variations at the soma seen in Fig. 3a top row are solely due to the realistic irregular soma shapes, which are quite different from simplified spherical/ellipsoidal approximations. After that, all faces within other neurons have been removed (Section 2.3).

**Fig. 3.**
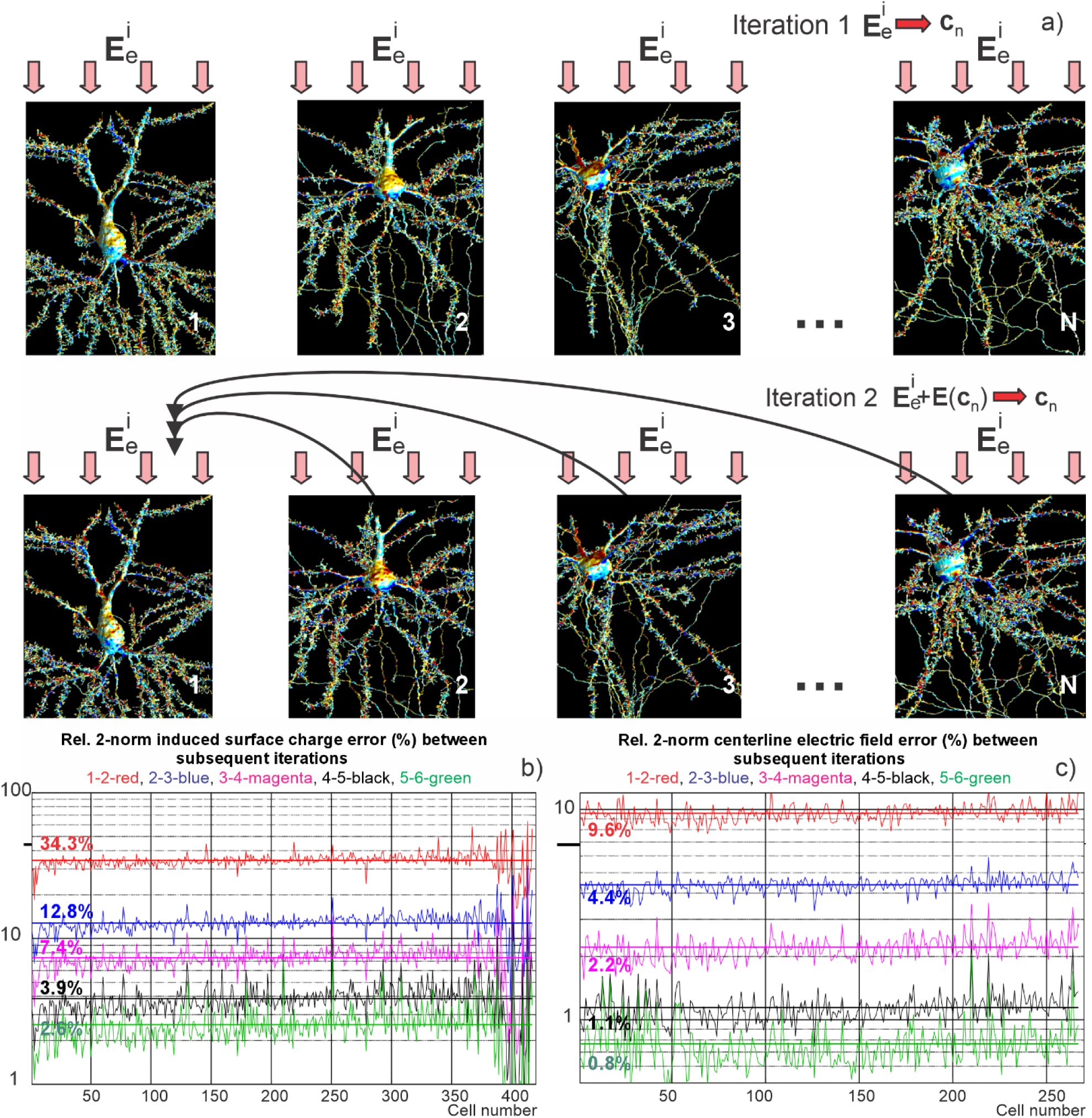
a) Nested iterative solution method. At step 1, every neuron is considered independent. Membrane charge deposition in response to impressed or primary field ***E***^***i***^ (***E***_**e**_^***i***^ in the extracellular space) is found via an iterative BEM-FMM solution. At step 2, the primary field ***E***_**e**_^***i***^ is modified by the field of charges deposited on all other neurons. Membrane charge deposition for every neuron in response to this modified field is found next. The process further repeats itself until a convergence criterion is met. b) Convergence curves for the relative 2-norm surface charge density error. c) Convergence curves for the relative 2-norm error for the collinear electric field projected onto centerlines of neuronal processes.

At step 2, the primary field ***E***^***i***^(***E***_**e**_^***i***^ in the extracellular domain) is modified by the field of charges deposited on all other neurons found in step 1. The membrane charge deposition for every neuron is recomputed with this modified primary field. At the following steps, the process repeats itself until a convergence criterion is met. In Fig. 3, the convergence criterion used required the relative 2-norm induced surface charge density error to be below 3% on average and the relative 2-norm centerline electric field error to be below 1% on average.

The corresponding convergence curves (Fig. 3b,c) indicate a sub-percent error for the collinear electric field at the neuronal centerlines after 6 iterations. An additional test of solution accuracy has been performed by comparing with the global BEM-FMM solution for 11 neighbor neurons treated as a single network. This test is described in Supplement E. There, the relative average charge density error for every neuron versus the global iterative or ‘exact’ solution for that single network after six iterations was approximately 1%.

Emphasize that the neuron-to-neuron interactions are taken into account at every step of the nested iterative solution except trivial first iteration (Fig. 3a). In its turn, the nested iterative algorithm used in this study provides a complete solution; it rapidly converges to the exact result when the number of iterations becomes sufficient (as shown in Supplement E).

The BEM-FMM solution is already highly parallelized, so we did not attempt additional parallelization of the nested iterative loop. An outer loop selects a neuron under study. An inner loop retrieves the charges on all other neurons found previously and computes their fields on the facets of the neuron under study (adds these fields up). Next, the neuron under study is solved, i.e., its new charge distribution is obtained, etc.

This method is potentially capable of modeling an unlimited number of cells, although CPU time now increases as *O*(*N*^2^) where *N* is the total number of cells. Electromagnetic computational results of this study required approximately 3 months of continuous CPU time when simultaneously utilizing multiple terminal servers and nodes of a high-performance computing cluster.

Fig. 4 illustrates the final induced charge map for the dorsal-ventral (*y*) polarization. It demonstrates the induced surface charge density for neuron #321 and its 200 nearest neighbors. Fig. 4a,b is the induced surface charge density for neuron #321 in sagittal and coronal planes. Fig. 4a is the induced charge density when only neuron #321 is present (iteration 1 or self-interaction). Fig. 4b is the induced charge density when all neurons and microcapillaries of the sample are present (iteration 6). Fig. 4c,d is the induced surface charge density for 200 nearest neighbors of neuron #321. Fig. 4c gives the data for the complete network of 200 nearest neighbors. Fig. 4d is a zoomed in area (x33) of the network in Fig. 4c with some inter-synaptic interconnections clearly visible. We were unable to plot electromagnetic data for more than 200 neurons simultaneously due to the limitations of the graphical software used.

**Fig. 4.**
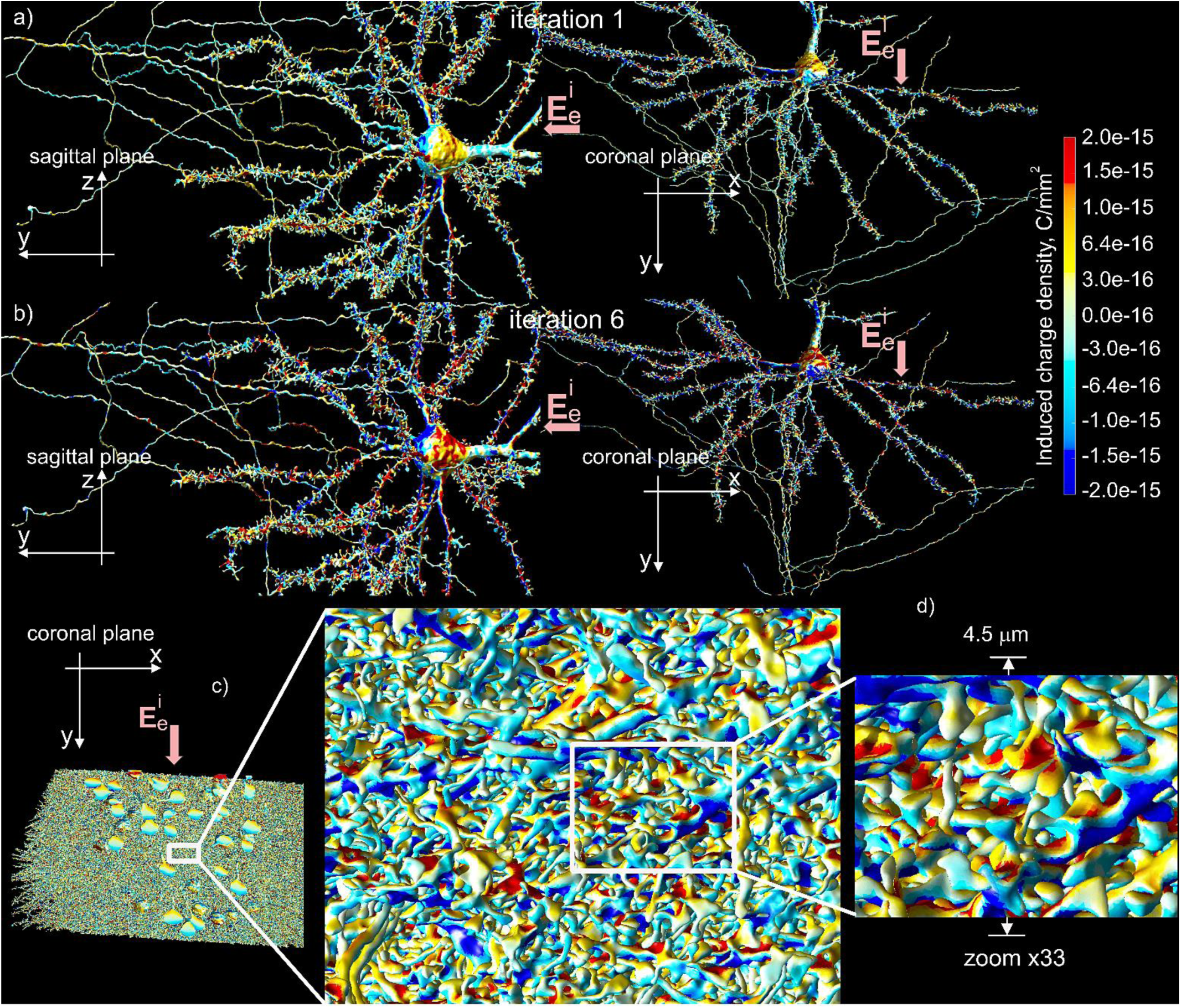
a,b) Induced surface charge density for neuron #321. a) Induced charge density when only neuron #321 is present (iteration 1 or self-interaction). b) Induced charge density when all neurons and microcapillaries of the sample are present (iteration 6). c,d) Induced surface charge density for 200 nearest neighbors of neuron #321. c) Complete network of 200 nearest neighbors. d) Zoomed in area (x33) of the network in c). We were unable to simultaneously plot electromagnetic data for more than 200 neurons due to the limitations of the graphical software used.

### 2.6 Average extracellular potential computed at centerlines/skeletons of neuronal processes

The activating function of the one-dimensional cable equation [41],[42],[43] needs extracellular potential or the collinear extracellular electric field at the centerlines of neuronal processes in the form of infinitesimally thin, transparent ‘wires’. For finite-thickness neuronal processes, these values are extracellular fields averaged over the cross-section. In the extracellular domain (*and only there*), the field solution obtained previously is equivalent to the solution for the entirely non-conducting interior. Therefore, the averaged values are most accurately obtained when the field and the potential are directly computed at the centerlines using the same extracellular solution which simultaneously holds for the entirely non-conducting interior [32],[33].

Fig. 5 illustrates the present averaging method (Fig. 5b) and the resulting effect of membrane charge deposition on the collinear electric field projected onto the centerlines of neuronal processes for neuron #321 (Fig. 5a,c). The sagittal plane view is shown corresponding to Fig. 4a,b left. Fig. 5a is the electric field projection in homogeneous space, when the neuronal membrane is not present per se and the same unform impressed electric field of 100 V/m is applied along the *y* axis (from dorsal to ventral). Fig. 5c is the averaged collinear electric field projection when all neuron membranes are present, and the primary field is distorted by the field of the induced charges. One observes that perturbations of the collinear electric field are quite significant. Moreover, they are constructively or “in phase” accumulated in space so that a positive perturbation is added to the originally positive field projection and vice versa.

**Fig. 5.**
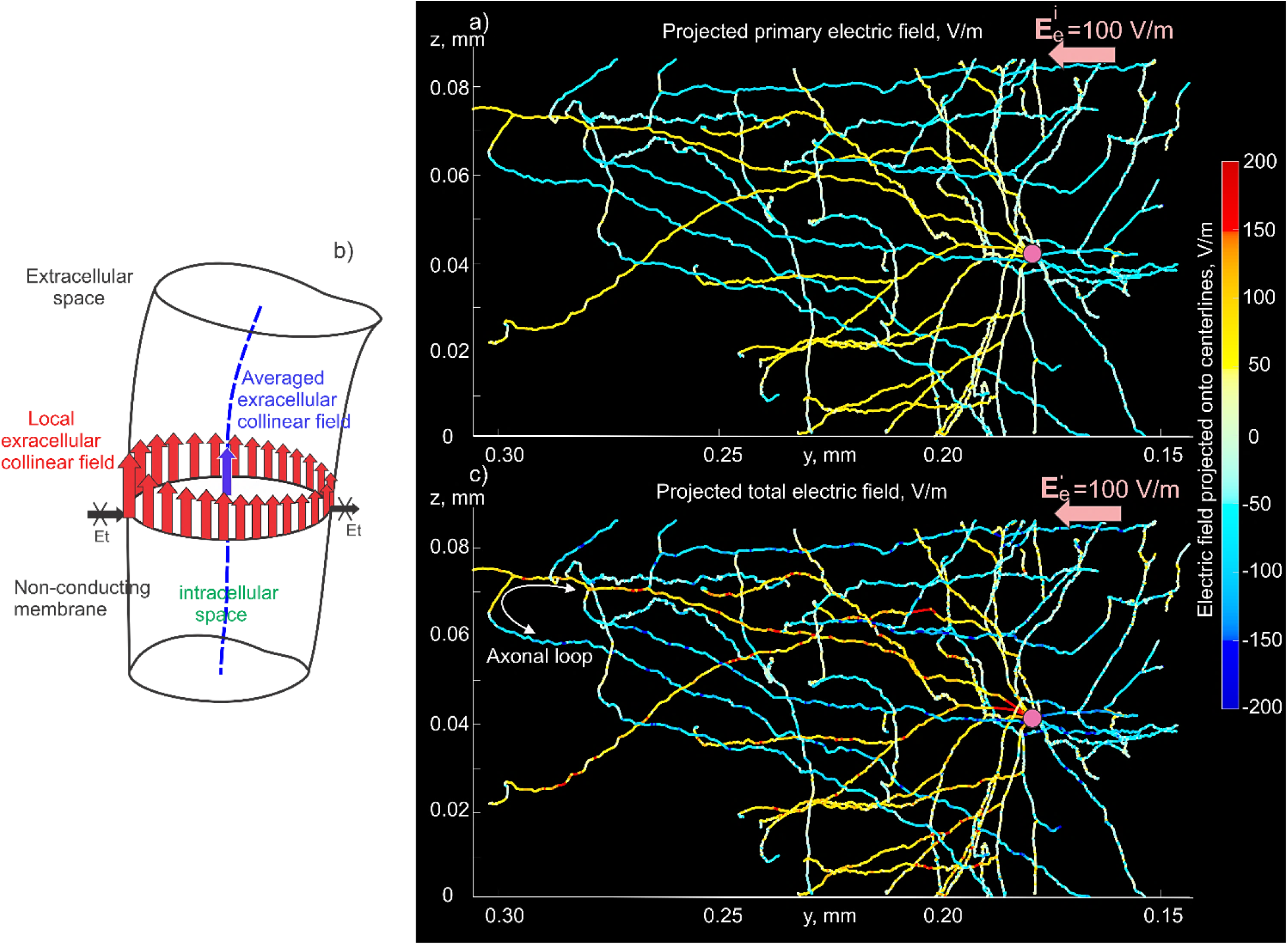
Effect of membrane charge deposition on the collinear electric field projected onto the centerlines of neuronal processes for neuron #321. The sagittal plane view is shown corresponding to Fig. 4a,b. a) Electric field projection in homogeneous space, when the neuronal arbor is not present and a uniform impressed electric field of 100 V/m is applied along the *y* axis (from dorsal to ventral). b) Averaging method over a cross-section. c) Electric field projection when all neurons are present, and the primary field is distorted by the field of all induced charges.

An appealing result of this approach is that both the transmembrane potential and the double layer density at the end of initial polarization are obtained automatically. Once the extracellular potential *φ*_e_ is known, we subtract its value from the constant resting potential value inside the cell and obtain the transmembrane potential *φ*_m_. Its product with the electrical permittivity of the membrane is the density of the double layer of charges across the cell membrane.

### 2.7 Biophysical modeling

We implemented rodent neuron models developed in Refs. [46],[10],[11],[12] with 6 active channels for the soma, 8 active channels for the axons, 1 active channel for the basal dendrites, and 3 active channels for the apical dendrites initially developed and used by Aberra et al. [10]. The NEURON software [45] with fine spatial and temporal resolutions has been employed to study neural activation in response to the extracellular potential/electric field computed previously at the centerlines of the neuronal processes. The neuron morphology for 267 available neuron models (including soma, axons, apical and basal dendrites) was loaded into NEURON by its Import3D tool. The implementation for the most important axonal channels was additionally verified against in-house MATLAB-based biophysical software. Every neuron was modeled independently, following the approach of Refs. [10],[11],[12]. Intercellular communications were not yet considered.

We used four stimulation pulse forms (Fig. S3 of Supplement C): (i) a monophasic positive/negative rectangular pulse starting at *t* = 0 and with a duration of 0.1 ms and; (ii) an experimentally measured positive/negative biphasic TMS pulse with an overall duration of 0.3 ms. The pulse amplitude gradually increased until the neuron activating criterion (the transmembrane voltage rises above zero anywhere in the arbor) was met. After that, the amplitude was decreased with a smaller step to pass the activating threshold in the opposite direction. This automated search procedure – the bisection method – requires approximately 15 steps and about 15-20 min per neuron on average. The action potentials were partially initiated at the axonal terminations following the previous observations [10],[11],[12]. For some neurons – primarily where the neuronal processes have been substantially cut out – it was also the axon hillock.

The overall simulation time for all neurons with *x*, *y*, and *z* polarizations, for all pulse forms, and for both primary and total extracellular fields/potentials did not exceed two CPU months. Other details of the biophysical modeling setup are described in Supplement C.

The L2/L3 dataset does not provide information about axon myelination. However, the rodent axons are partially myelinated [10],[11],[12]. To account for this, we also modeled the case of artificial myelination. All axonal branches were assigned a myelin sheath with a thickness of 0.2 times the local radius. The nodes of Ranvier were placed at bifurcations, terminations, and along the segmental length, 20 µm apart from each other.

### 2.8 Correction of average extracellular field due to finite sample size

Replacing a block of homogeneous tissue with the microscopic realistic specimen in a conventional macroscopic brain model must not change the solution in the surrounding homogeneous tissue. This in particular means that the extracellular conductivity of the heterogeneous specimen must be higher than the surrounding homogeneous conductivity, to enable nearly distortion free current flow through the specimen with non-conducting membranes, which are partially blocking the extracellular current flow. To do so, a box around the specimen is needed. For the present specimen (the non-orthogonal parallelepiped in Fig. 1c), it is difficult to put it in a tightly enclosing rectangular box without substantial (up to 25%) reduction of the already small cellular volume, closing hundreds of holes, etc.

Therefore, an artificial rectangular cuboid sample mimicking the original specimen was constructed as shown in Fig. 6. Fig. 6a shows an equivalent straight fiber density distribution (computed in Supplement A) of the sample to be converted to the form of a filled rectangular cuboid. The fiber spacing there schematically illustrates weak sample anisotropy. Fig. 6b is an artificial computational equivalent of the specimen from Fig. 6a constructed using ∼500 finite cylinders in the form of a 3D lattice, with the overall mesh size of ca 2 million facets. The cylinders are spaced equally, but they have proportionally different radii to account for the moderate sample anisotropy from Fig. 6a. The cylinders are embedded into an enclosing (yellow) box (250×140×90 µm) whose conductivity contrast (interior vs. exterior conductivity) can be arbitrarily varied. The volumetric fill factor of the cylindrical lattice within the box is 0.3, which coincides with the fill factor for the original sample (Supplement A). The average extracellular field within the box is evaluated for 0.7 million uniformly distributed observation points. Cylinder walls (i.e., ‘membranes’) and their tips, which are protruding the box as I Fig. 6b, are assumed to be non-conducting. The field within the cylinders with the non-conducting walls (in the ‘intracellular’ space) is not computed.

**Fig. 6.**
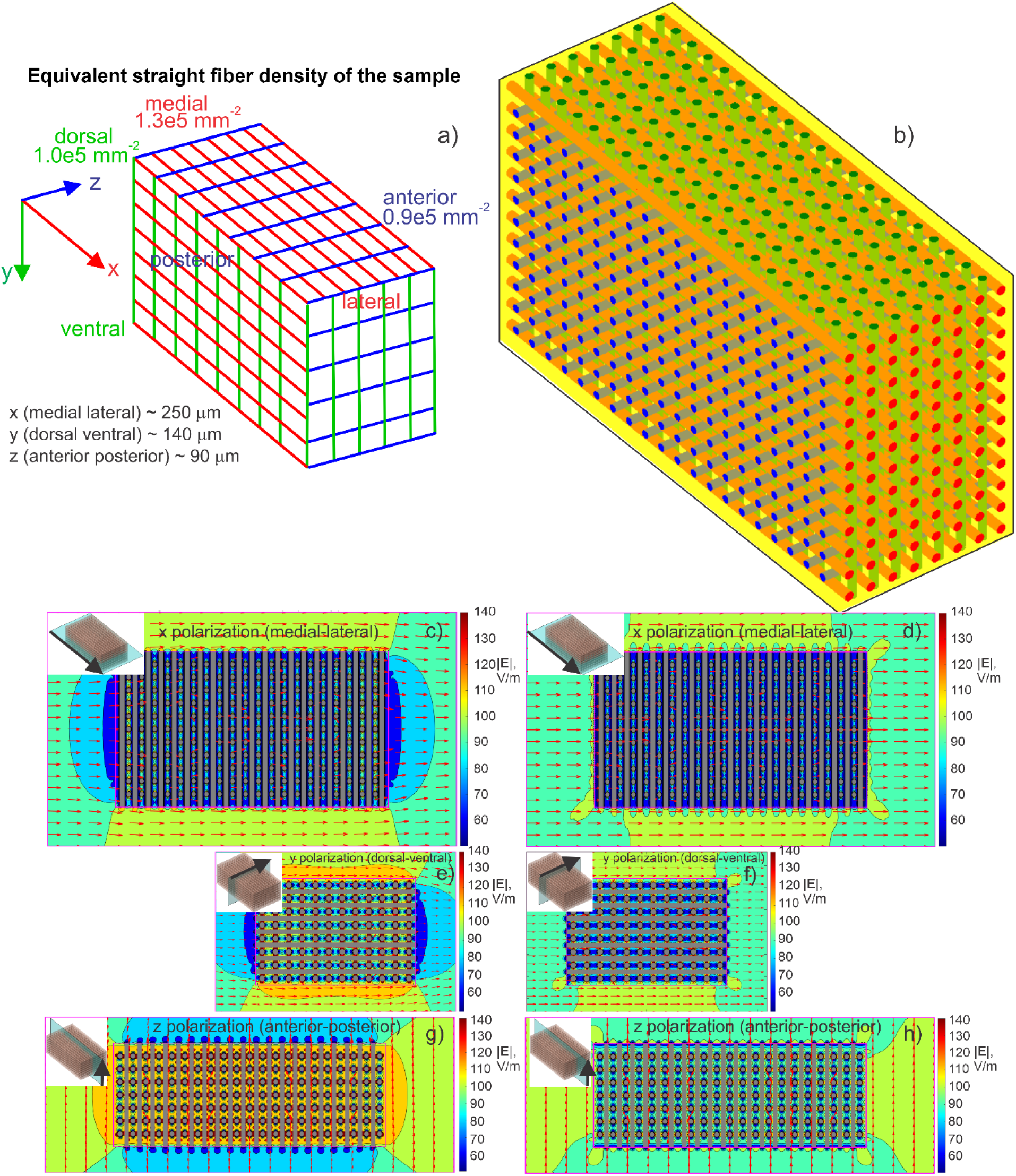
a) Equivalent straight fiber density distribution of the sample converted to the form of a rectangular cuboid. The fiber spacing illustrates weak sample anisotropy. b) Artificial computational equivalent of the specimen from a) constructed using ∼500 finite cylinders in the form of a 3D lattice. The cylinders are spaced equally but have proportionally different radii to account for the moderate sample anisotropy from a). The cylinders are embedded into an enclosing (yellow) box (250×140×90 µm) whose conductivity contrast (interior vs. exterior conductivity) is varied. The volumetric fill factor of the cylindrical lattice within the box is 0.3, which coincides with the value for the original sample. Cylinder walls (membranes) and their tips, which are protruding the box, are assumed to be non-conducting. c-h) Magnitude (in V/m) and direction of the total extracellular electric field within and around the artificial sample for three different polarizations (*x*, *y*, *z*) of the impressed field ***E***^***i***^ with the value of 100 V/m. c,e,g) – Field distortion around the heterogeneous artificial sample when its extracellular conductivity is that of the surrounding homogeneous medium. d,f,h) – Field distortion around the heterogeneous artificial sample when its extracellular conductivity is increased with an attempt to achieve a minimum surrounding distortion.

Fig. 6c-h shows the magnitude (in V/m) and direction of the total extracellular electric field within and around the artificial sample for three different polarizations (*x*, *y*, *z*) of the impressed field ***E***^***i***^with a value of 100 V/m. Fig. 6c,e,g illustrates the field distortion around the heterogeneous artificial sample when its extracellular conductivity is that of the surrounding homogeneous medium. Fig. 6d,f,h provides the field distortion around the heterogeneous artificial sample when its extracellular conductivity is increased with an attempt to achieve a minimum overall distortion in the surrounding space.

The external field distortion was estimated in three orthogonal planes shown in Fig. 6c-h. The extracellular conductivity was varied with the goal of minimizing the distortion over the outer plane area for all three planes simultaneously. An optimal extracellular conductivity increase by a factor of 1.7 was found using brute force optimization. Note that this conductivity increase within the sample appears to be close to that predicted by the Maxwell equation for dilute cell suspensions [57],[58],[59].

The correction factor is the ratio of the average extracellular field magnitude within the sample with the original and increased conductivity, respectively. This ratio was found to be equal to 1.08 for the *x*-polarization, 1.14 for the *y*-polarization, and 1.23 for the *z*-polarization. The activating thresholds found in the original specimen were multiplied by those values to account for sample matching to the surrounding homogeneous space.

## 3. Results

### 3.1 Neuronal activating thresholds within the sample vs homogeneous medium

We chose the first 90 neurons with the longest non-cut axonal arbors exceeding 0.5 mm (Supplement A). The neurons have been renumbered in descending order based on the length of their axonal arbors. This operation eliminates, to the best of our abilities, neurons with severely truncated axonal arbors (at the sample boundaries) from the subsequent analysis, which would otherwise yield non-physical results. Fig. 7 shows the computed activating thresholds within the sample matched to the surrounding homogeneous medium compared to those in the microscopically homogeneous medium. Here and in the following, we only consider the 90 “best” neurons.

**Fig. 7.**
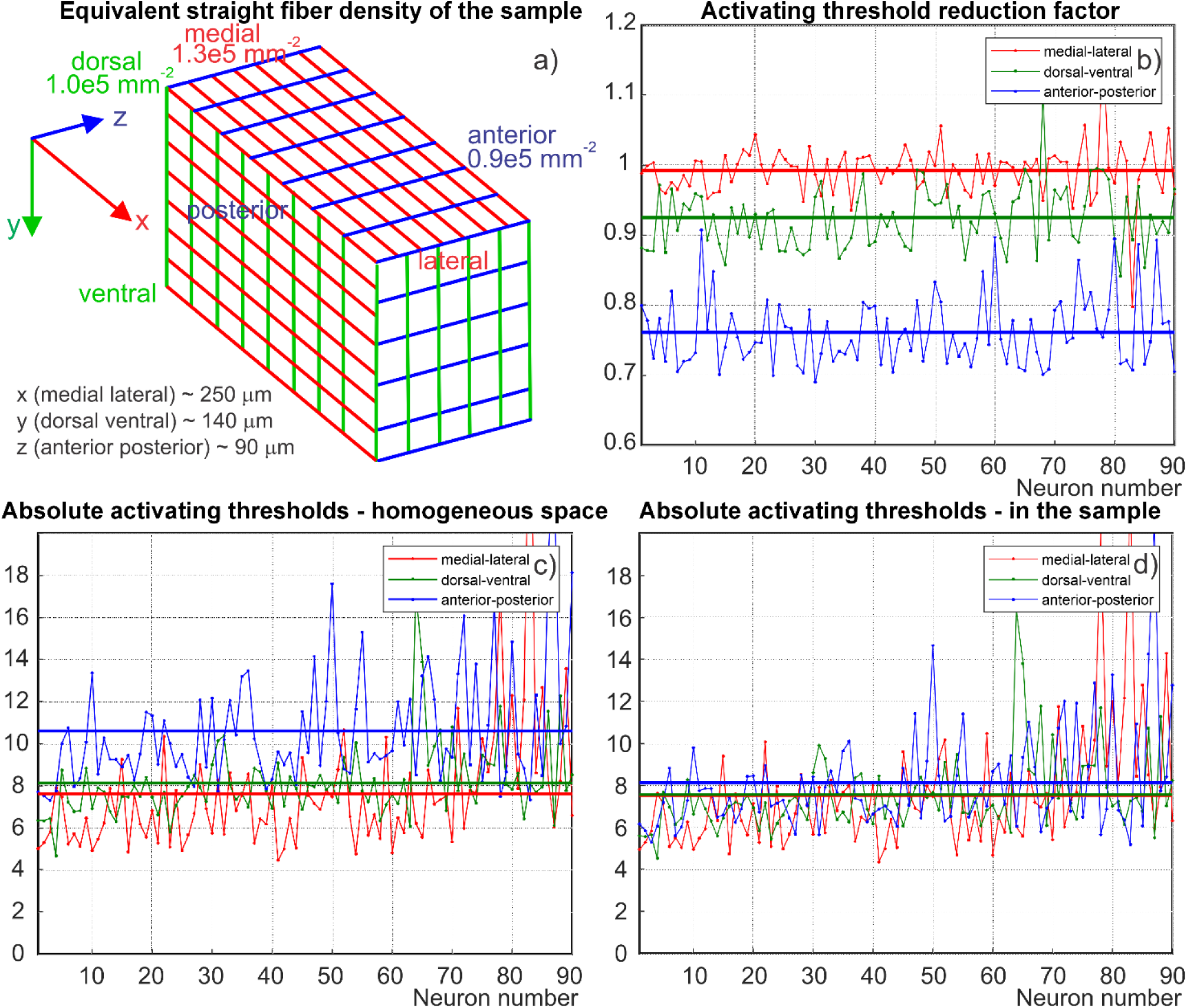
a) Equivalent straight fiber density distribution of the sample characterizing its anisotropy. b) Activating threshold reduction factor for the first 90 neurons with the longest axonal arbors (sorted) for three polarizations of the primary electric field: medial-lateral (red), dorsal-ventral (green), and anterior-posterior (blue). c) Absolute activating thresholds for the same 90 neurons in homogeneous space. d) Absolute activating thresholds for the same 90 neurons within the sample. The units in c,d) are given in terms of the base field strength of 100 V/m; one unit corresponds to 100 V/m. These results are given for the case of the unmyelinated axonal arbor and a negative rectangular activation monophasic pulse with the duration of 0.1 ms. Results for all other cases are given in Supplement C.

Fig. 7b gives a neuron-by-neuron threshold reduction factor. The reduction factor is defined here as the ratio of the activating threshold of a physical neuron within the sample (with the total extracellular field ***E***_e_ and the total potential *φ*_e_) to the activating threshold of the neuron skeleton in the homogeneous medium (subject to the field ***E***^***i***^ and the potential *φ*^***i***^). The threshold reduction results in Fig. 7b are given for three polarizations of the primary electric field: medial-lateral (red), dorsal-ventral (green), and anterior-posterior (blue) and for the 0.1 ms long rectangular negative stimulating pulse.

Fig. 7c gives the corresponding absolute activating thresholds for 90 neuron skeletons in the homogeneous space. Fig. 7d gives the absolute activating thresholds for the same 90 physical neurons within the sample. The units in Fig. 7c,d are given in terms of the field strength of 100 V/m; one unit corresponds to 100 V/m. Figs. S4-S10 of Supplement C report the same data for the remaining pulse forms and for myelinated axons. The last figure there was computed for the positive biphasic 0.3 ms long pulse. The data in Supplement C are conceptually similar to those shown in Fig. 7. Average threshold reduction factors for all pulse forms and polarizations of ***E***^***i***^are summarized in Table 1.

**Table 1.** Averaged relative threshold reduction factor of the neuronal activating thresholds for different polarizations of the impressed (primary) electric field and different pulse forms (Supplement C) between a neuronal skeleton in a homogeneous medium and the physical neuron in the brain sample, respectively. The first 90 neurons with the longest (uncut) axonal arbors have been processed. Since data for myelinated axons are quite similar (Supplement C), only one such dataset has been included. Correction for the average field was found by modeling an artificial rectangular cuboid of the same effective size and fill factor including anisotropy from Fig. 6b (Section 2.8).

| Modeling setup | Medial-lateral (x)<br>polarization of $E^i$ | Dorsal-ventral (y)<br>polarization of $E^i$ | Anterior-posterior<br>(z) polarization of<br>$E^i$ |
| --- | --- | --- | --- |
| A Monophasic rect. pulse, 100 μs, negative | 0.99 | 0.93 | 0.82 |
| B Monophasic rect. pulse, 100 μs, positive | 0.97 | 0.97 | 0.83 |
| C Biphasic pulse (TMS), 300 μs, negative | 0.97 | 0.96 | 0.83 |
| D Biphasic pulse (TMS), 300 μs, positive | 0.96 | 0.97 | 0.83 |
| E Same as A but with all axons myelinated | 0.98 | 0.93 | 0.80 |
| Average | **0.97** | **0.95** | **0.82** |

### 3.2 Deviation of activating thresholds for different polarizations of primary field

Table 2 summarizes normalized standard deviations of the average neuronal activating thresholds for three different polarizations of the primary electric field. They are computed as std([*X Y Z*])/mean([*X Y Z*]) where *X Y Z* are the average thresholds for three respective polarizations. The first 90 neurons with the longest axonal arbor have again been processed. From Table 2 and Fig. 7, one observes that the normalized standard deviations between average thresholds for different directions of the applied primary field are always smaller for the sample as compared to individual neuron skeletons in homogeneous space. Cell membrane interactions apparently tend to make brain stimulation more ‘equal’ in all directions.

**Table 2.** Normalized standard deviation of the neuronal activating thresholds for three different polarizations of the primary electric field (from Fig. 7 and Supplement C). The first 90 neurons with the longest (largely uncut) axonal arbors have been processed.

| Modeling setup | Normalized STD:<br>Homogeneous medium | Normalized STD:<br>Microscopically realistic<br>brain |
| --- | --- | --- |
| A Monophasic rect. pulse, 100 μs, negative | 0.18 | 0.04 |
| B Monophasic rect. pulse, 100 μs, positive | 0.35 | 0.23 |
| C Biphasic pulse (TMS), 300 μs, negative | 0.18 | 0.12 |
| D Biphasic pulse (TMS), 300 μs, positive | 0.15 | 0.11 |
| E Same as A but with all axons myelinated | 0.22 | 0.19 |
| Average | **0.22** | **0.14** |

### 3.3 Strength-duration results

Fig. 8a illustrates a weak dependency of the threshold reduction factor on the duration of the applied stimulation pulse. A case in point is neuron #348 within the sample when the negative rectangular pulse is applied. At the same time, the absolute values of the activating threshold indeed follow the classic strength duration dependency [61] with a high degree of accuracy. This is shown in the inset in Fig. 8a; note that a log-log scale has been used there.

**Fig. 8.**
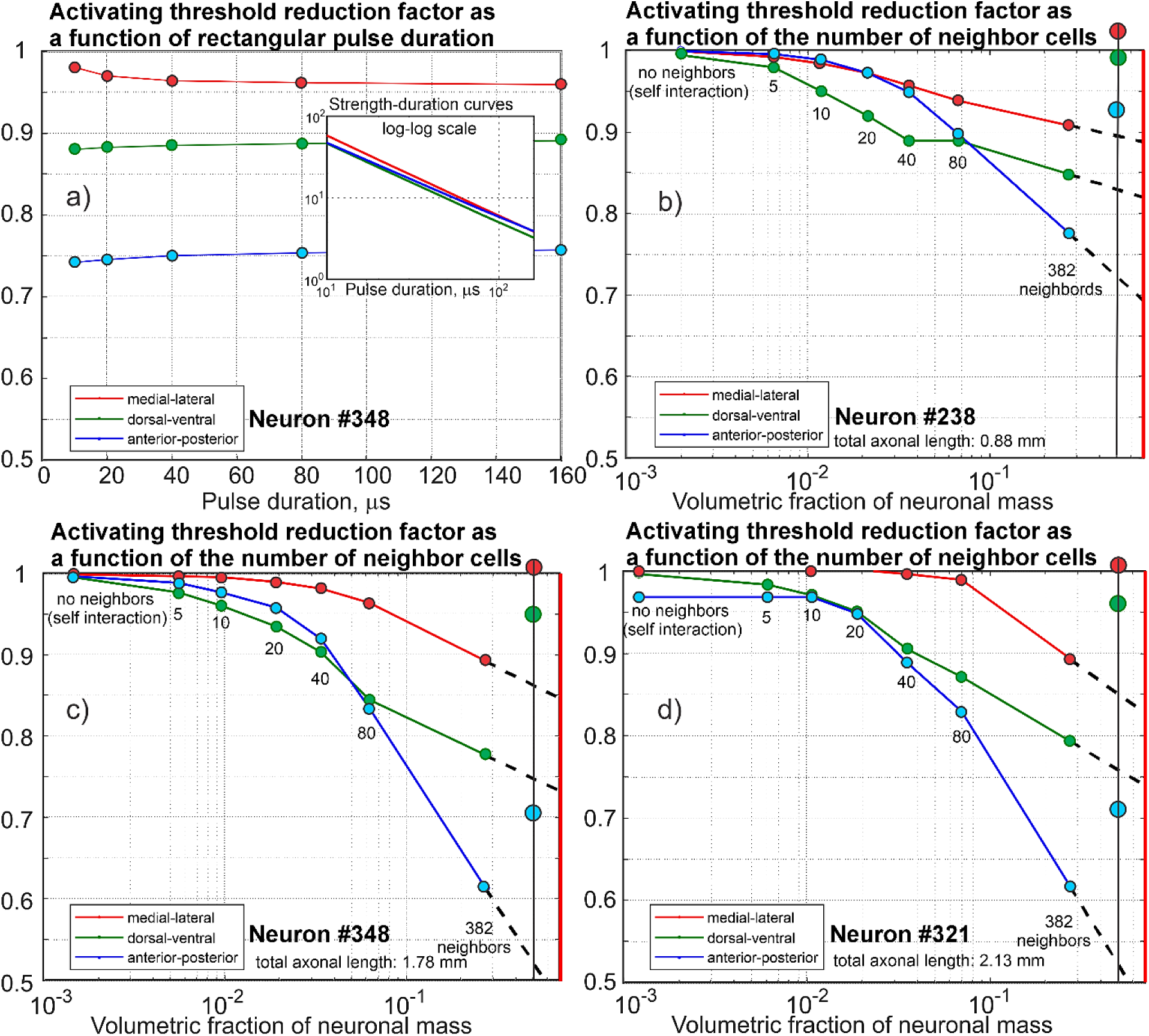
a) Activating threshold reduction factors as functions of rectangular negative pulse duration. The inset shows the corresponding strength-duration curves for neuron #348 within the brain sample. Inset units in are in terms of the initial field strength of 100 V/m; one unit corresponds to 100 V/m. b,c,d) Activating threshold reduction factors as functions of the number of neighboring cells for three representative excitatory neurons with different lengths of the axonal arbor (for the rectangular negative pulse) without the sample polarization correction. All three neurons belong to the set of 90 neurons with the longest axonal arbor. The x-axis is the corresponding volumetric fraction of neuronal mass computed separately for each neuron when sequentially adding its neighboring cells. Red lines show the expected fraction of 70% [50]; dashed lines show expected extrapolation results. Vertical black lines on the right and the corresponding larger circles are the results with the sample polarization correction for volumetric concentration of 0.5.

### 3.4 Dynamics of threshold reduction factor as a function of a number of surrounding neurons

Fig. 8b,c,d illustrates the threshold reduction factor dynamics as a function of the number of neighboring cells included in the computations for three representative neurons located close enough to the sample center. For each of three representative neuron cells (#238, 321, 348, all having different lengths of axonal arbor), we start adding its nearest neighbors (0, 5, 10 neighbors, etc.) from the entire brain sample and then repeat both electromagnetic and biophysical computations to find new activating thresholds. The *x*-axis in Fig. 8b,c,d is the corresponding relative volumetric fraction of neuronal mass computed using known volumes of each cell. The first point of every curve is the threshold reduction factor without neighbors, which is due to self-interaction of the neuron itself; the last point is the result for complete brain sample. These curves do not yet include the sample polarization correction, which was very difficult to obtain for the scattered neurons.

### 3.5 Extrapolation of threshold reduction factor to more realistic neuronal density

The volumetric fraction of the neuronal cells within the volume was estimated in Table S1 (Supplement A) as ∼30%. This is less than the known value of ∼60-70% [48],[49],[50] in rats and mice since all neurons with somas outside the brain sample were omitted and their dendritic/axonal processes, which are still within the sample, were ignored. The threshold reduction factor curves in Fig. 8b,c,d could in principle be extrapolated (as shown by dashed lines in Fig. 8b,c,d) to estimate the threshold reduction factor for the likely more realistic volumetric neuronal fill factor of 70% (shown by a red vertical line in Fig. 8b,c,d).

To estimate the effect of the sample polarization correction, we again computed the expected extracellular field decrease for one volumetric concentration of 0.5 following the method of Section 2.8. In Fig. 8b,c,d, the vertical black lines on the right and the larger colored circles are the final results with the sample polarization correction.

## 4. Discussion

### 4.1. Methodology

The present study is a method to electromagnetically model a realistic microscopic brain sample in application to brain stimulation. Our target region is a relatively small L2/L3 mouse primary visual cortex brain sample (250 ×140×90 μm^3^ volume) or IARPA Phase I [5],[6],[7] with less than 400 tightly packed neurons, with much of the neuronal arbor missing due to excluding cells with somas located outside the volume under study, and with an underdeveloped microvasculature.

We show that the boundary element fast multipole method or BEM-FMM [27],[28],[29] – when using its charge-based formulation [30],[31],[32],[33] – is an appropriate tool for the combined electromagnetic and biophysical analysis of not only this sample, but also potentially more accurate modern detailed brain samples [7],[8],[9] with the resolution of at least 100 nm under certain assumptions. These assumptions include:

i. Only major scatterers or perturbers of the applied electric field – neurons themselves and microcapillaries – are included into consideration. The remaining cells are combined into one extracellular space. This assumption may be revised with regard to astrocytes and oligodendrocytes [9].
ii. Cell membranes are assumed to be nonconducting at the end of the initial polarization period for a duration of several microseconds [34]. This assumption remains valid until the end of a stimulation pulse (with a duration of approximately 0.1-1 ms). In this time interval, we compute the activating function via the one dimensional cable equation [41],[42],[43] by averaging the total collinear extracellular field over a cross-section of a neuronal process.
iii. The neuronal cells then experience a change in their physiological state with the characteristic time scale of several milliseconds or longer [34]. At this point, we are again following the one-dimensional cable equation of the standard biophysical theory but with the activating function equal to zero.

Our modeling method is not limited to the present case of a uniform applied brain stimulation field, which is more common for transcranial magnetic and electrical stimulation. It can equally well target any type of brain stimulation, including biophysical aspects of deep brain stimulation [66],[67],[68] and intracortical microsimulation/brain-computer interfaces (cf., for example [69],[70],[71]).

### 4.2. Lowering activating thresholds?

Our initial hypothesis was that microscopic extracellular electric field noise might substantially ease neuron activation compared to macroscopically homogeneous brain modeling. However, this hypothesis was generally not confirmed by the present modeling results. More specifically:

- The average activating thresholds remain nearly the same for electric field polarizations aligned along the two longest sample directions – medial-lateral and dorsal-ventral (see Table 1).
- Some minor variations (reduction factors) observed there are primarily attributed to lowering of activating thresholds for neurons with already high absolute activating thresholds. Such neurons do not significantly contribute to recruitment. On the other hand, the neurons with the lowest activating thresholds – the most sensitive neurons (the ones of real interest) – are mostly unaffected (cf. Fig. 7 and the corresponding figures of Supplement C).
- Although the average activating threshold for the anterior-posterior direction of the applied field does become lower, it is again significantly affected by lowering activating thresholds for neurons with already high absolute activating thresholds (Fig. 7 and the corresponding figures of Supplement C). Furthermore, this is the shortest sample dimension. There, the absolute activating thresholds are high and the effect of sample polarization might not be entirely undone.
- Extrapolation to a more realistic neuronal mass yields a similar result (Fig. 8).

Still, the fiber in the anterior-posterior direction is subject to the largest number of intersections with perpendicular fibers per unit length, as evidenced by Fig. 7a, and is proportional to the fiber density in the two other directions, i.e. to 1.0*e*5 × 1.3*e*5. For fibers in the dorsal-ventral direction, the number of such intersections per unit length in Fig. 7a is medium; it is again proportional to the fiber density in two remaining directions, i.e. to 0.9*e*5 × 1.3*e*5. Finally, for fibers in the medial-lateral direction, the number of such intersections per unit length in Fig. 7a is the lowest and is proportional to the fiber density in two other directions, i.e. to 0.9*e*5 × 1.0*e*5. These three numbers of intersections per unit length relate as 1.0, 0.90, and 0.70 and correlate well (Pearson correlation coefficient of 0.999) with the activating threshold reduction factors from Fig. 7b. Therefore, further work with larger regularly shaped samples may be necessary to prove or disprove this effect and estimate its strength.

### 4.3. When might field perturbations be important?

Consider, for example, a simple numerical microsimulation experiment with a Hodgkin-Huxley axon originally studied by Rattay [43]. A cathodic stimulation of the 10 mm long axon by a point electrode located at the origin is shown in Fig. 9a. A monophasic electric pulse (*T* = 0.1 ms) is applied. Field perturbers are 2D cylinders with non-conducting walls located perpendicular to the figure plane and with a large radius of 200 µm. Charge deposition on the cylinder walls perturbs the electrode field from Fig. 9a as shown in Fig. 9b where the total field is displayed. As a result, an electric field ‘spatial noise’ with a mean value of zero appears for the extracellular collinear electric field along the axon, as shown in Fig. 9c in blue. The noise amplitude is large; it can approach or even exceed the applied field itself when the scatterers are in close proximity to the axon (cf. basic solutions for DC current conduction, for example from [47]).

**Fig. 9.**
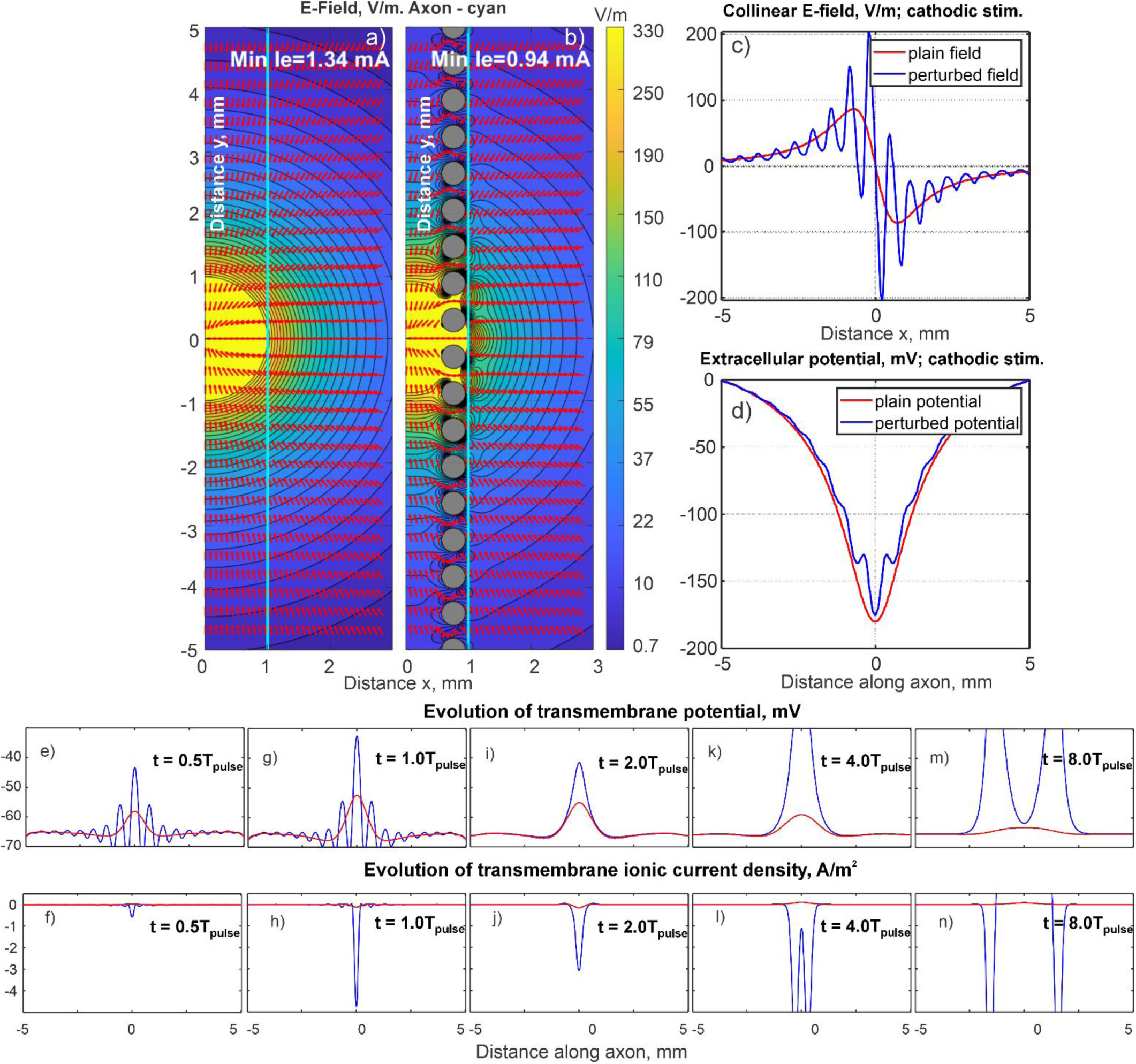
Concept of the electric field spatial noise. a) Straight HH axon from Ref. [43] and the electric field of cathodic stimulation. b) The same case as in a) but with artificial field perturbers – infinite cylinders. c,d) Collinear electric field and extracellular potential with (blue) and without (red) field perturbations. In both cases, the electrode current is 0.94 mA. e-n) Resulting evolution of transmembrane potential and transmembrane ionic current density for both cases at different times. Red curves are those without field perturbations; blue curves correspond to the perturbed extracellular field.

When *t* < *T* (Fig. 9e,f), the transmembrane potential closely follows the activating function (the derivative of the collinear electric field) with sharp positive peaks of the latter indicating membrane depolarization [43]. The inward (negative in Fig. 9) ionic current flows within the depolarization peaks (Fig. 1f) while nothing flows within the hyperpolarization peaks – the noise is being ‘rectified’. This process is strongly nonlinear – cf. Fig. 9f,h. There, one local inward current injection starts to dominate other inward injections while all outward ionic currents are still negligibly small. Despite the following low pass filtering as in Fig. 9i,j, this local current injection appears to be strong enough to support further action potential generation (Fig. 9k,l,m,n) when the stimulation cathodic current is only 0.91 mA. This contrasts with the effect of the unperturbed field from Fig. 9a which would generate an action potential if the cathode current were 1.31 mA or higher. Thus, the threshold reduction factor due to the spatial noise is 0.91/1.31=0.69 in the present case. When the cylinder radius in Fig. 9b is halved (becomes 100 µm), the threshold reduction factor increases to 0.84. When the radius is further halved and is reduced to 50 µm, the threshold factor becomes 0.98 and then quickly approaches one when this process continues. Similar results hold for the mouse axon.

It appears that strong yet very spatially narrow field perturbations caused by smaller cylinders do not have a significant effect, as they diffuse quickly along the cable with no lasting impact. This observation is directly evident during the simulations and is valid at the terminals as well. According to this example, to have an effect, the inward current injections must be not only strong but also sufficiently extended in space. The product of strength times the spatial length appears to be crucial. Proper lengths are observed in the case of blood vessels or other “unexpected” scatterers, such as possibly a combination of 2-3 nearby somas, a large astrocyte or oligodendrocyte, a favorable combination of several smaller scatterers, etc.

According to the present simple example, the reduction effect becomes apparent only for relatively large scatterer sizes, on the order of ∼0.1λ where λ is the axonal space constant. This constant is equal to 1.5 mm for the Hodgkin-Huxley axon and to 0.6 mm for the mouse axon studied here. A similar observation holds for a finite axon (either Hodgkin-Huxley [43] or mouse [10],[11],[12]) in a uniform field subject to a pair of cylindrical scatterers at its termination or to a bandlimited electric field noise applied to an entire axon. The simple MATLAB code used to create this example is described in Supplement F and made available online [72].

### 4.4 Limitations of this study

The small size of the L2/L3 brain sample leads to cutting out a large portion of the neuronal processes, most notably axon collaterals. As an example, the average length of the existing axonal arbor per neuron is ∼0.5 mm for the present L2/L3 sample, ∼2.5 mm (with the maximum length of 36 mm!) for the 1mm^3 sample from Allen Inst. [7],[8], ∼6 mm for Markram’s *et al* elongated data in L2/L3, and ∼7 mm for Markram’s *et al* data in L5 [46].

Our computations show that the absolute activating threshold values are severely affected by the total axonal length. Additionally, the arguably most sensitive excitatory neurons of L5 [11] are not included into the sample. This is perhaps why the absolute threshold values in Fig. 7 and Figs. S4-S10 (Supplement C) overestimate the experimentally measured value of ∼90 V/m in mouse primary motor cortex (Supplement D) very significantly. Even in the most favorable case of myelinated axons, there are only nine neurons with the activating threshold below 200 V/m (Fig. S8, Supplement C) and none of them has the activating threshold below 100 V/m.

Furthermore, the blood network of the L2/L3 sample was not the major target of IARPA Phase I study. It is therefore severely underdeveloped (cf. Fig. 1d) compared to more recent and accurate results [7],[8],[9].

Our biophysical modeling treats all neuronal cells independently, following previous work [10],[11],[12] to demonstrate the impact of realistic microscopic structures on direct neuron activation threshold. On the other hand, the L2/L3 brain sample is already supplied with a list of ∼2,000 synapses between 334 excitatory pyramidal cells [7] obtained in [6] allowing future extension of our approach to simulating network/synaptic modulation [64]. Furthermore, we only considered the uniform impressed stimulation field and a single-pulse stimulation time course.

Though used ubiquitously, there are inherent limitations of the one-dimensional cable equation, especially close to the large somas. Modern bidomain models (cf. [44]) could possibly resolve this issue.

The next logical yet extremely challenging step is to analyze the much larger 1mm^3 MICrONS mouse brain sample, which includes a considerably better-developed axonal arbor and ∼75,000 neuronal cells, and another very recently published human brain sample [9]. This would require further improvements and modifications of the computational method.

## 5. Conclusions

A method is proposed to accurately model perturbations of an impressed extracellular electric field within a microscopically realistic brain, with many tightly spaced neuronal cells. This impressed field could be due either to brain stimulation or to a firing cluster of neurons themselves. For the first time, we have been able to create a map of extracellular electric field distributions within the relatively sizable fully anatomical L2/L3 mouse brain sample (250 × 140 × 90 μm³) containing ∼400 detailed neuronal cells and some blood microcapillaries with the resolution of 100 nm.

Electric field modeling was further coupled with 1D biophysical modeling, which is based on extracellular field averaging, neuron morphologies precisely linked to the physical neurons and known ion channel models for rodents. This combination was applied to predict changes in neuron activating thresholds caused by microenvironment – the electric field spatial noise with a mean value of zero on the cellular level.

Under stated assumptions and limitations, our results show that the microscopic field perturbations – the electric field spatial noise caused by the neuronal structure – only modestly influence the predicted stimulation field strengths necessary for neuronal activation. This result supports the conventional theory of “invisible” neurons embedded in a macroscopic brain model [10],[11],[12], for transcranial magnetic and transcranial electrical stimulation. Although large in magnitude, the microscopic activating function perturbations are predominantly limited to very narrow spatial regions (e.g., dendritic diameters) so that the local inward current injections diffuse rather quickly instead of triggering substantial changes in the physiological state. However, our result is sample-specific and subject to a number of limitations and restrictions as stated above.

## Acknowledgements

Authors express their gratitude to Dr. Bethanny Danskin and the Virtual Observatory of the Cortex (VORTEX) program of the Allen Institute for Brain Science for their assistance in processing the L2/L3 brain sample, to Dr. Clayton S. Bingham of NINDS/NIH and Dr. Cameron C. McIntyre of Duke University for shaping early stages of this study, to Dr. Ted Carnevale of Yale School of Medicine for his guidance in utilizing NEURON software with superior spatial resolution, and to Dr. Padma Sundaram of Massachusetts General Hospital for useful discussions.

Special thanks are extended to Dr. Ermal Toto and Dr. James Kingsley of Academic and Research Computing at Worcester Polytechnic Institute for their great support in assembling and executing highly demanding parallel computational tasks. This study has received support from the NIMH grant R01MH130490 and NIBIB grant R01EB035484 (SNM), NIDCD grant R01DC020891 (ARN), NINDS grant U24NS120053 (BD) as well as the Intramural Research Program of NIH, NIMH ZIAMH002955 (Z-DD), and Intramural Research Program of NIH, NIDA ZIADA000638 (HL).

Finally, we express our sincerest thanks to the two anonymous reviewers for their dedication and comprehensive, professional comments, which shaped both the methods and conclusions of this study. We are also thankful to the deputy editor for their effort.

## Supplement A Post-processing segmentation and skeletonization of L2/L3 sample

### A1. Preparing neuronal surface meshes for electromagnetic modeling

All 396 neurons were initially numbered following the y-coordinate (dorsal-to-ventral direction) of their center of mass. The lowest numbers correspond to the deepest neurons where the bulk of the axonal arbor was cut out (see below). Neurons with the highest numbers have the (relatively) well developed axonal arbor (Fig. 1 of the main text).

Initial computational surface triangular meshes for the neurons downloaded from the MICrONS database [1] were not entirely suitable for accurate numerical analysis. Automated CGAL alpha wrapping C++ library [2],[3] was used for all neurons to re-mesh the data and convert them to the strictly 2 manifold high-quality (min triangle quality is 0.1) high-resolution individual surface objects. A relative α (vs 1e-3*longest diagonal; parameter α limits the maximum triangle size) of 1000 and a relative offset (vs 1e-3*longest diagonal, parameter offset limits the tightness of the result around the input mesh) of 4000 were used in the CGAL program. We were unable to properly heal 13 cells. These cells were eliminated from the analysis which reduced the total cell number from 396 to 383.

Fig. 2 of the main text illustrates the final membrane mesh for neuron #238. A typical membrane mesh has approximately 1-2 million triangles on average, with an average facet size kept at 100 nm. This is necessary to keep a proper spatial accuracy for intersection resolution (Section 2.3 of the main text) and also resolve microscopic, localized variations of the impressed electric field. The total number of facets of the sample approaches 0.5 billion. The average facet size is 100 nm. An identical procedure also was applied to the microcapillaries (cf. Fig. 1d of the main text).

### A2. Volumetric density of neuronal cells computed from surface meshes and its comparison with literature values

The volumetric fraction of all neuronal cells within the SEM (scanning electron microscopy) volume was estimated as 30%±5% by computing the volume of each neuron using the CGAL tools and also taking into account neuronal mesh intersections (Section 2.3 of the main text).

This is less than the known estimates in rats and mice [4],[5],[6] because all neurons with somas outside the brain sample were omitted and their dendritic/axonal processes, which are still within the sample, were ignored.

In general, the brain tissue is composed of extracellular space and intracellular space, i.e. neurons and glia cells (intracellular space) do not take up all the volume of brain tissue. Oxygen molecules and nutrients (e.g. glucose molecules) diffuse from capillary through extracellular space to neurons and glia, and are taken up through transporters. According to [4], neuronal elements occupy about 74% and glial elements about 8% of the total volume in rats. According to [5] (rats), the extracellular space takes up about 20% of total cortex volume in adult rats. According to [6] (Table 5), the fraction of neuronal mass in mice cerebral cortex is 70%.

Based on these data and also on some data from Ref. [7], we estimate the realistic volume fraction of neuronal mass in mice neocortex as approximately 70%.

### A3. Healing and refining centerlines of neuronal processes – neuron skeletons

The original neuronal skeletons or morphological reconstructions (301 in total) from the MICrONS database [1] didn’t remain entirely in the center of the corresponding neuronal processes represented by the relevant surface mesh. They also had a rather sparse discretization. The skeleton branches or centerlines have been therefore refined, smoothed, and corrected via an automated in-house routine that finds every point’s distance from the enclosing membrane walls in the up, down, left, right, in, and out directions and averages them for every cross-section of the relevant triangular mesh. Fig. 2b,c of the main text shows the centerlines of the neuronal processes healed in such a way. We attempted to maintain a nearly uniform spatial resolution of 100 nm for the centerline discretization(observation) points, which should be sufficient for catching up fast field variations over the length on the order of 1 µm or so. We were unable to properly heal 34 centerlines. These centerlines were eliminated from the subsequent analysis which reduced their total number from 301 to 267.

We also estimated the average axonal length per neuron, with the maximum length of 8.3 mm for inhibitory neuron #191 and the minimum length of 0 mm (for 39 other neurons in total). The latter case is non-physical; it is a deficiency of the centerline extraction method.

### A4. Estimating sample anisotropy from neuron morphology

To estimate the degree of anisotropy, we introduced an equivalent straight–fiber density measure. It is obtained when centerlines for all neuronal processes including both axons and dendrites are projected onto three Cartesian axes (Fig. S1b,c,d below). This density was estimated by finding the total projection length and then dividing this value by the product of three sample dimensions (the sample volume). The following values of the equivalent straight fiber density have been obtained: 1.3*e*5 mm^−2^ for the medial-lateral or x direction, 1.0*e*5 mm^−2^ for the dorsal-ventral or y direction and 0.9*e*5 mm^−2^ for the anterior-posterior or z direction.

### A5. Selecting proper neurons with the longest retained axonal arbors for biophysical modeling

The lengths of all segments of the neuronal processes (axons and dendrites) were computed using the available `*.swc` files from the MICrONS database. Fig. S1a shows the length of axonal arbor per neuron, now sorted in descending order. It is seen that neurons with numbers greater than 90-100 have a vanishingly short axonal arbor, which was cut out of this sample almost entirely. Therefore, they were eliminated from the final results of biophysical analysis (otherwise non-physical, large values of the activating threshold could be obtained) but were still kept for the electromagnetic analysis.

Thus, only the neurons with the overall axonal length of 0.5 mm or greater (with the sorted numbers from 1 to 90, Fig. S1a below) were included into the relevant biophysical modeling outcomes – see Fig. 6 of the main text, Figs. S4 through S7 of Supplement C, and also Tables 1 and 2 of the main text.

Fig. S1b additionally shows the total length of axonal arbor per neuron projected onto three Cartesian axes (x, y, z). The neurons have been sorted as in Fig. S1a). Similarly, Fig. S1c demonstrates projected (onto three Cartesian axes) length of dendritic arbor per neuron. The neurons have again been sorted as in Fig. S1a. Finally, Fig. S1d demonstrates the projected length of all neuronal processes (axonal plus dendritic) per neuron. The neurons have been sorted as in Fig. S1a. Again, one clearly observes in Fig. S1b that the neurons with numbers greater than approximately 90 have vanishingly small axonal arbor, which was cut out of the brain sample almost entirely.

**Fig. S1.**
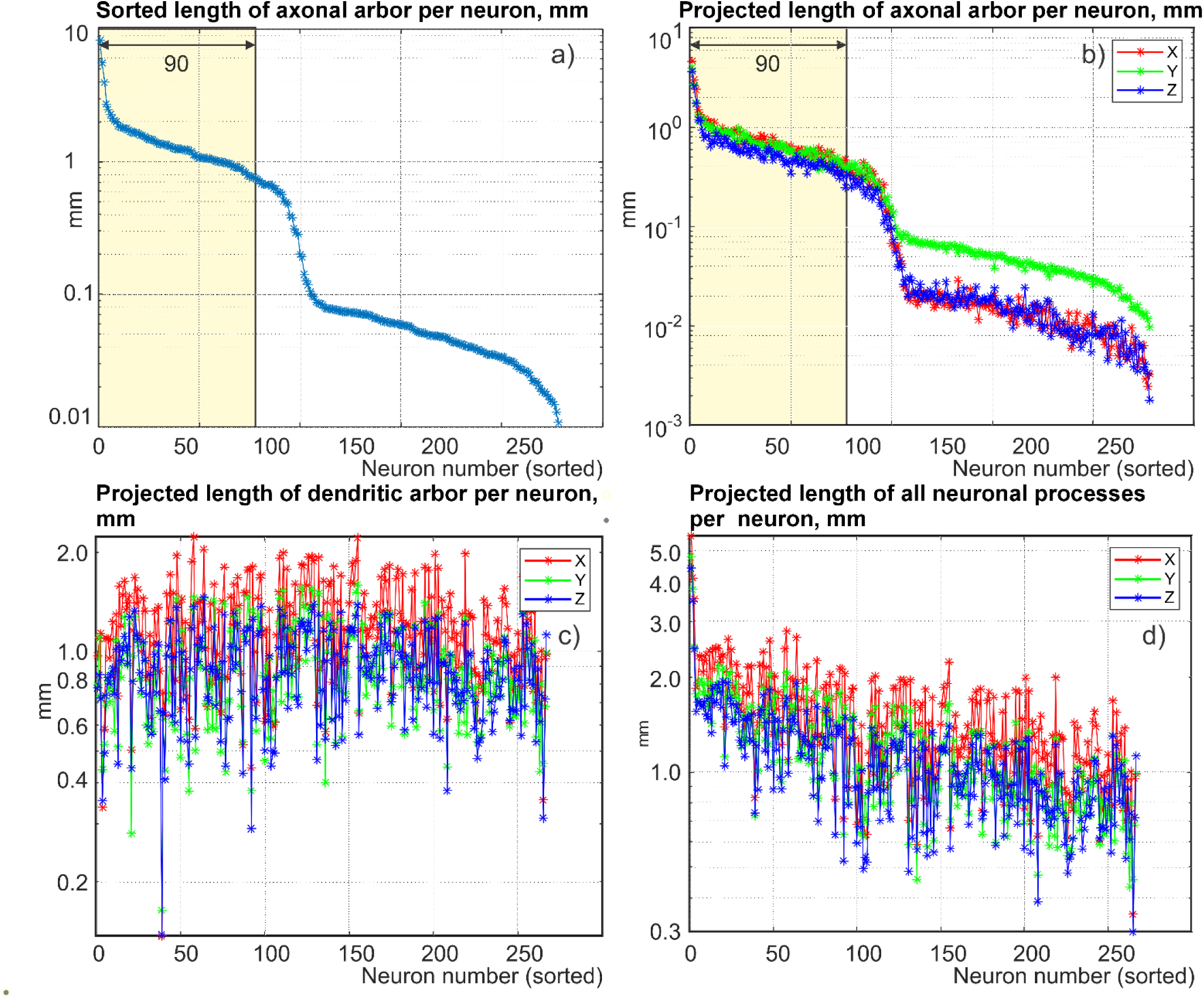
a) Sorted (in descending order) length of axonal arbor per neuron. Neurons with numbers greater than approximately 90 have vanishingly small axonal arbor, which was cut out of the sample almost entirely. b) Projected (onto three Cartesian axes) length of axonal arbor per neuron. The neurons have been sorted as in a). c) Projected (onto three Cartesian axes) length of dendritic arbor per neuron. The neurons have been sorted as in a). c,d) Projected (onto three Cartesian axes) length of all neuronal processes (axonal plus dendritic) for all neurons with the healed morphological reconstructions. The neurons have been sorted as in a).

### A6. Summary of post-processing parameters

Table S1 of this Supplement lists major topological and biophysical parameters of the created computational testbed (electromagnetic and biophysical) including estimated sample anisotropy. Emphasize again that both the facet size for membrane discretization and the skeleton’s spatial resolution for the biophysical analysis were kept at approximately 100 nm to resolve large yet very localized variations of the impressed field due to multiple nearfield interactions of the neuronal processes as well as to resolve multiple intersections of neuronal arbors of different neurons (Fig. 2 of the main text).

**Table S1.** Post-processing parameters of the L2/L3 mouse brain sample.383 Neurons were kept for electromagnetic analysis. Data for 90 neurons with the longest axonal arbor were included in the final biophysical outcomes.

| Parameter | Value |
| --- | --- |
| Neuronal meshes for EM modeling | 396 – initial; 383 – final |
| Vascular meshes for EM modeling | 34, partially disconnected |
| Total mesh size/triangular facet size/triangular facet quality | ~0.5 billion facets/100 nm/0.1 min |
| Volumetric fraction of neuronal mass within the sample | ~30±5% |
| Neuron skeletons | 301 initial/267 final |
| Spatial discretization of healed skeletons | 100 nm |
| Largest length of axonal arbor per neuron | 8.3 mm for inhibitory neuron #191 and 2.3 mm for excitatory neuron #333 |
| Equivalent straight-fiber density of neuronal processes in three orthogonal directions (sample anisotropy) | Fiber density of $1.3e5 \text{ mm}^{-2}$ for the medial-lateral or x direction, $1.0e5 \text{ mm}^{-2}$ for the dorsal-ventral or y direction and $0.9e5 \text{ mm}^{-2}$ for the anterior-posterior or z direction |
| Neuron skeletons with developed axonal arbor suitable for biophysical modeling | 90 |

## Supplement B Membrane mesh intersection maps

### B1. Mesh intersection maps

Fig. S2a demonstrates a typical surface mesh intersection map. Two neighboring excitatory neurons – #238 and #290 – have been chosen.

**Fig. S2.**
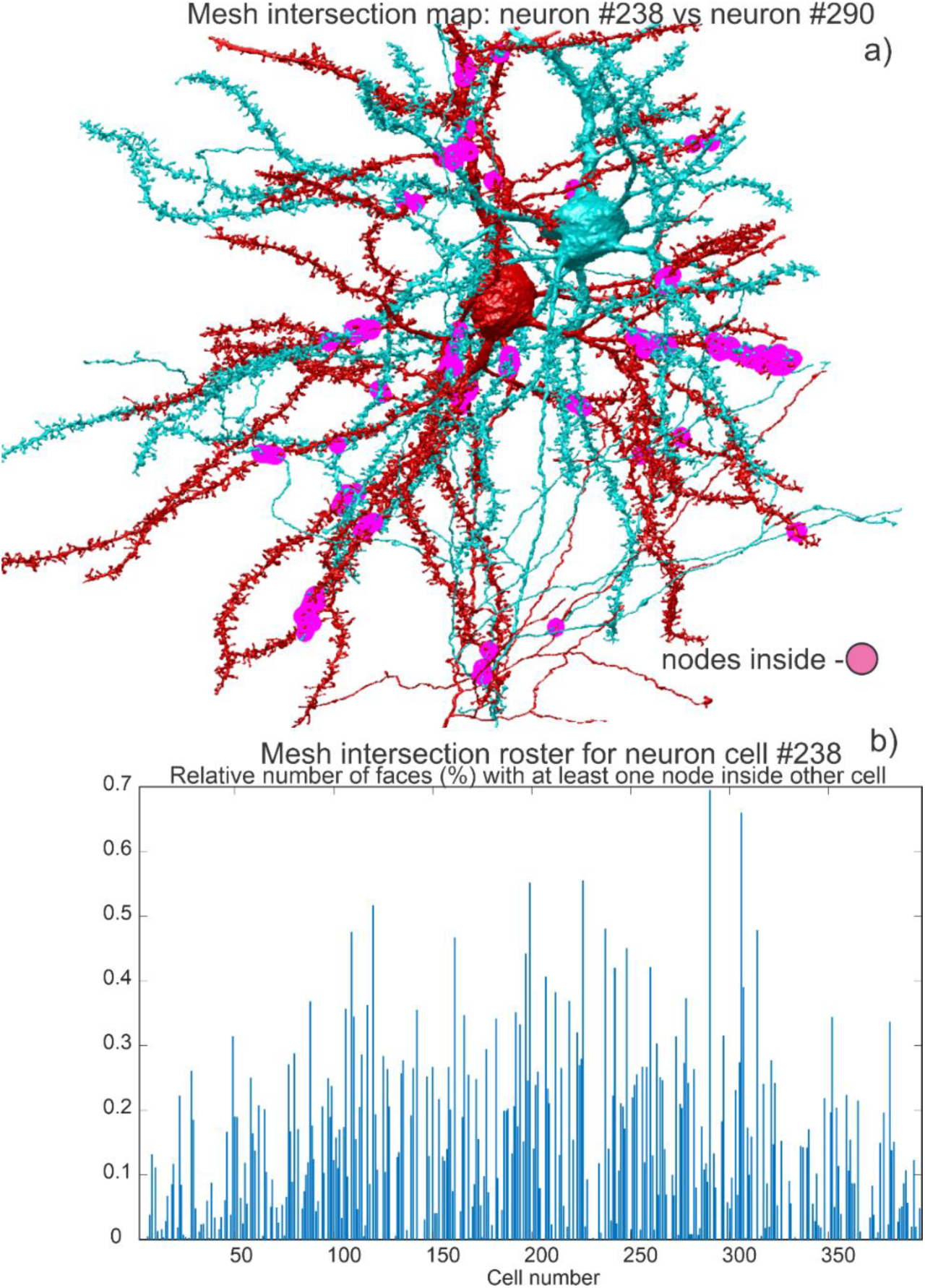
a) Surface mesh intersection map between neighboring excitatory neurons #238 and #290, respectively. Intersection domains are marked by magenta color. b) Mesh intersection roster of neuron #238 with all other neurons of the sample. The y axis gives the relative number of facets (% versus the total number of faces for neuron #238) with at least one facet node contained within another neuronal shell. The x axis is the neuron number.

In Fig. S2a, the mesh intersection domains are marked by magenta color. There are approximately 30 such independent intersection domains.

Fig. S2b also presents a mesh intersection roster of excitatory neuron #238 with all other neurons of the sample. It is observed in Fig. S2b that neuron #238 intersects with almost all neurons of the sample. Similar, and even more dramatic results were observed for other larger neurons of the sample. To identify the mesh intersections for large yet manifold meshes, we used vectorized function `inpolyhedron` written by Sven Holcombe in MATLAB [8]. Still, this has been a very time-consuming operation.

## Supplement C Biophysical modeling

### C1. Summary of NEURON modeling parameters

Standard NEURON software [9] has been used to study neural activation in response to the impressed extracellular potential/electric field computed previously at the centerlines of the neuronal processes. The post-processed neuron morphology (described in Supplement A) for 267 available neuron models (in `swc` format, including soma, axons, apical and basal dendrites) was loaded into NEURON by its `Import3D` tool. Following references [10],[11],[12] we applied the existing ion channel models for rodents (rats) which have been used starting with Aberra et al. [10]. We used 6 active channels for the soma, 8 active channels for the axons, 1 active channel for the basal dendrites, and 3 active channels for the apical dendrites. The implementation for the most important axonal channels was tested and verified against in-house MATLAB-based software.

The centerlines of the neuronal processes were discretized in space with the step *d_λ_* = *L*/[*n*_*seg*_*λ_w_*(*f* = 100 *Hz*)], where *L* is the section length, *n*_*seg*_ is the number of segments in this section, and 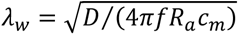 is the length constant, with *D* being the neuron diameter; *f* is a frequency that is high enough for transmembrane current to be primarily capacitive, but is still within the frequency range relevant to neuronal function. Here, *R_a_* is the resistivity of axoplasm and *c_m_* is the membrane capacity per unit area. In the computations, we tested several *d*_*λ*_ values between *d*_*λ*_ = 0.01 and *d*_*λ*_ = 0.0005, respectively, to address expected rapid field variations at microscale. We compared the results for a representative number (6 in total) of large neurons with many local extracellular field variations. The difference in the activating thresholds was found to be negligibly small between the two extreme values. Therefore, the value *d*_*λ*_ = 0.01 or 0.005 has been routinely used everywhere after test runs. The time step size was chosen as 1 μs which satisfies the stability condition.

### C2. Pulse forms of the stimulation field

As the simple base form, we used the monophasic rectangular pulse form for the primary or impressed field starting at *t* = 0 and with a duration of 0.1 ms – cf. Fig. S3a,b). The positive and negative pulses (positive and negative electric field directions) have been modeled separately. Along with this, we used a realistic biphasic TMS (transcranial magnetic stimulation) pulse form (Fig. S3c,d) from a B65 coil of MagVenture, Denmark measured in the Noninvasive Neuromodulation Unit, Experimental Therapeutics & Pathophysiology Branch, NIMH. Both positive and negative pulse forms have been investigated. Some computations have also been done for a monophasic TMS pulse.

The pulse magnitude was gradually increased until the neuron activating criterion (the transmembrane voltage goes above zero anywhere in the arbor [10]) was met. After that, the amplitude was decreased with a smaller step to pass the activating threshold in the opposite direction. This automated search procedure – the bisection method – requires approximately 15 steps and about 15-25 min per neuron on average. The total simulation time for all neurons with x, y, and z polarizations for both primary and total extracellular fields/potentials did not exceed two weeks per polarization.

**Fig. S3.**
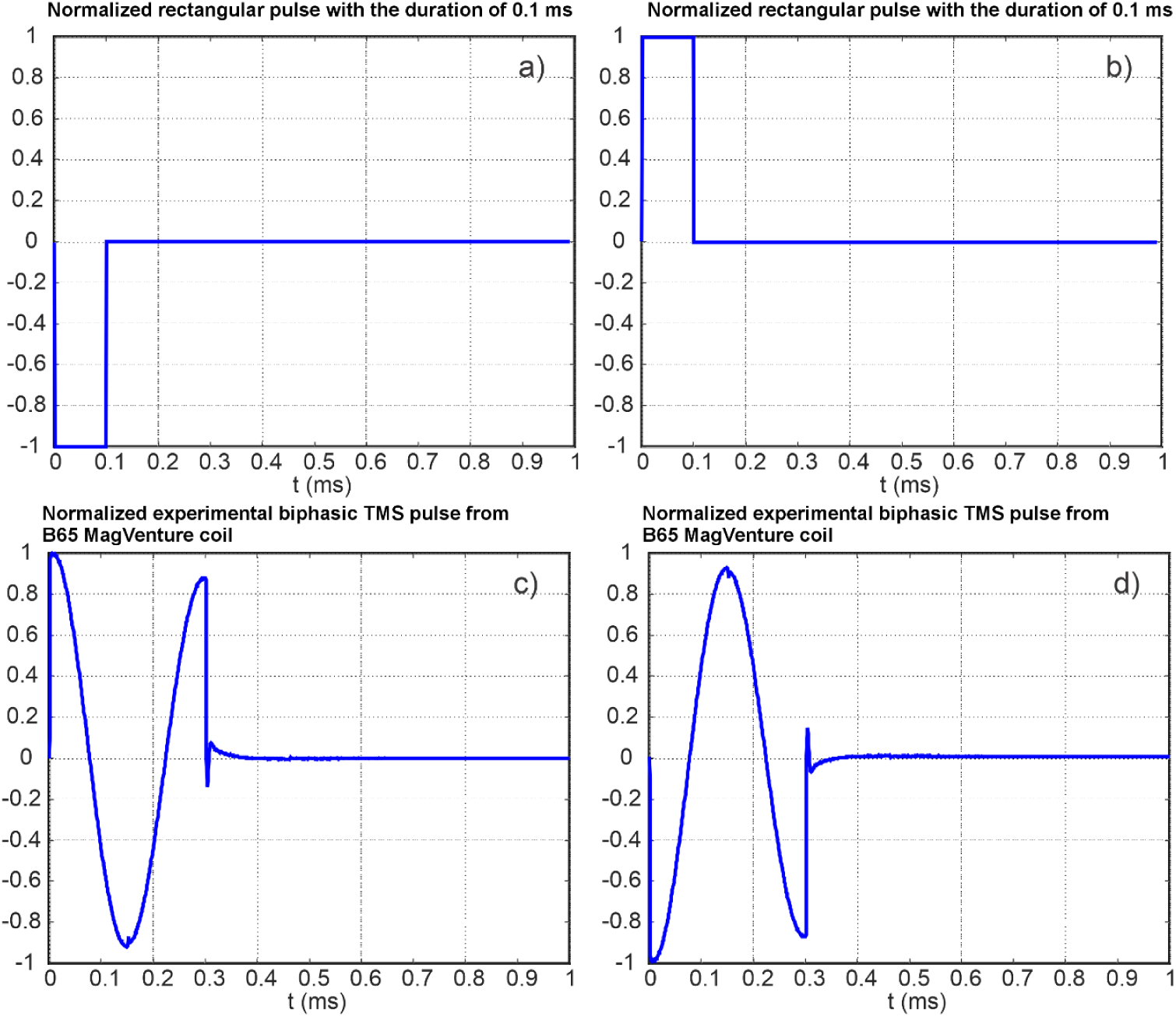
Four pulse forms for the primary impressed field used for biophysical modeling.

### C3. Reduction of neuronal activating thresholds for different pulse forms and myelinated axons

Figs. S4 through S10 given below replicate Fig. 7 of the main text for different pulse forms and with considering axonal myelination. They have been used to fill out Tables 1 and 2 of the main text.

**Fig. S4.**
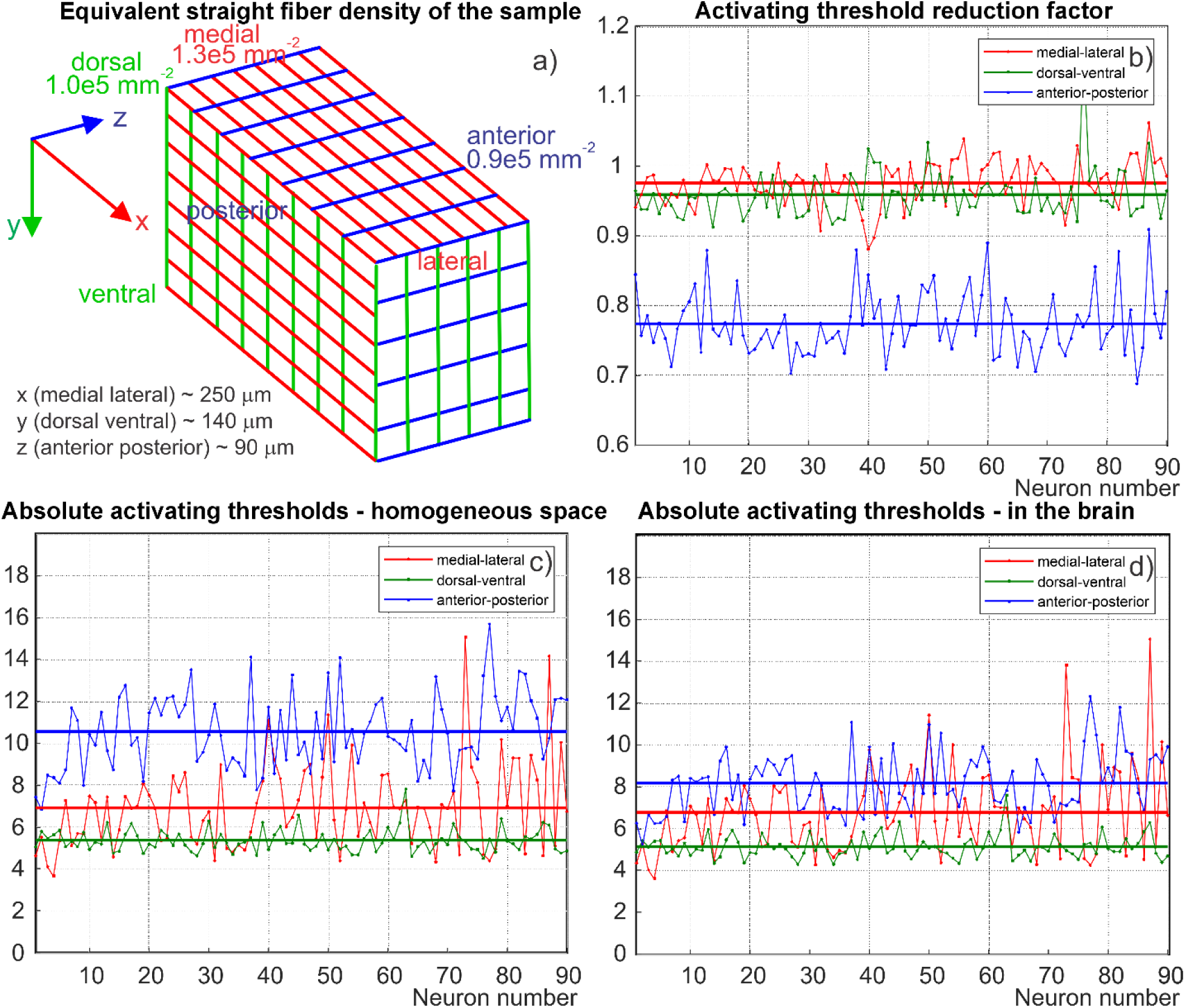
a) Equivalent straight fiber density distribution of the sample characterizing its anisotropy. b) Activating threshold reduction factor for the first 90 neurons with the longest axonal arbor (sorted) for three polarizations of the primary electric field: medial-lateral (red), dorsal-ventral (green), and anterior-posterior (blue). c) Absolute activating thresholds for the same 90 neurons in homogeneous space. d) Absolute activating thresholds for the same 90 neurons within the sample. The units in c,d) are given in terms of the base field strength of 100 V/m; one unit corresponds to 100 V/m. These results are given for the case of the unmyelinated axonal arbor and a positive rectangular activation monophasic pulse with the duration of 0.1 ms.

**Fig. S5.**
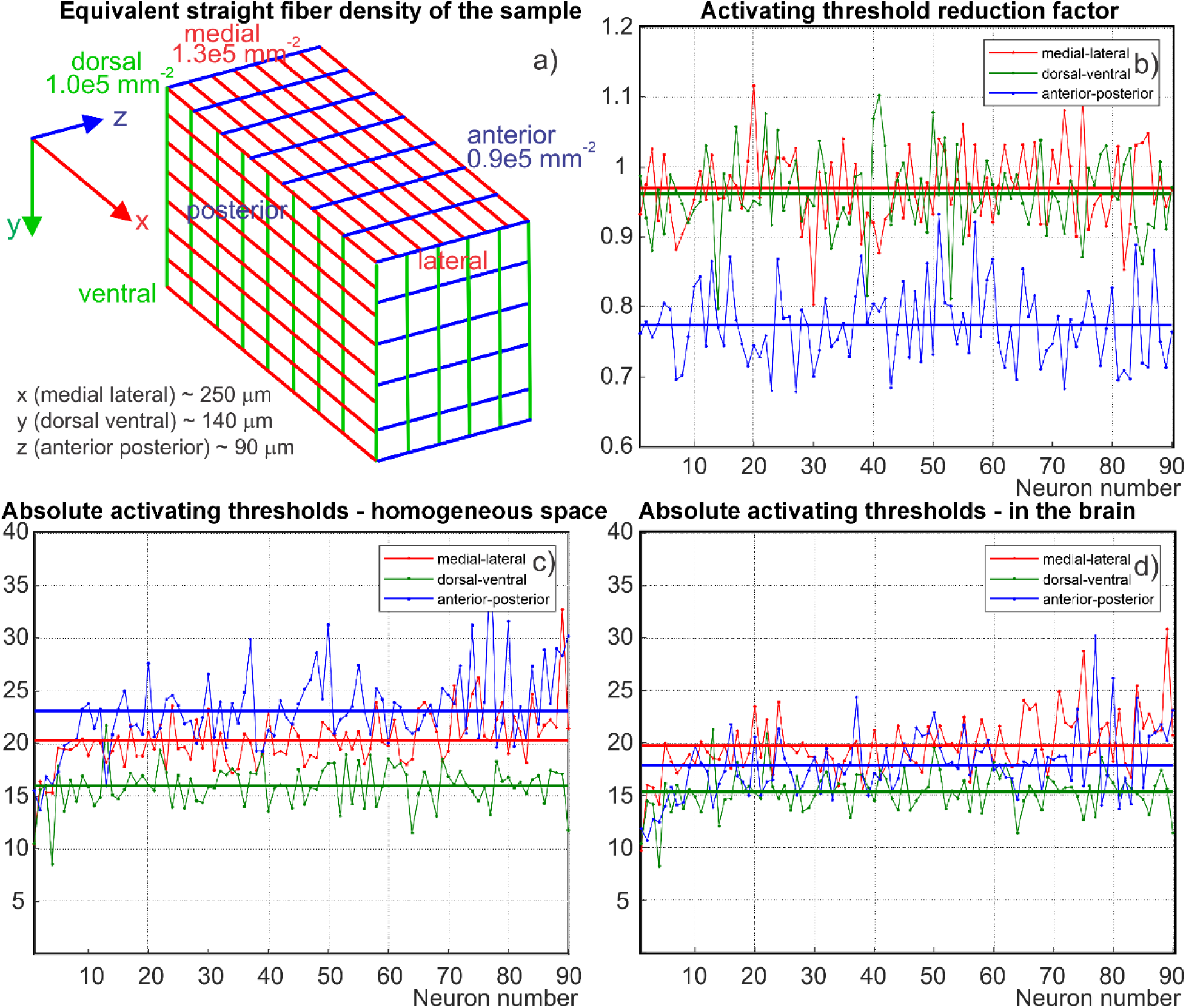
a) Equivalent straight fiber density distribution of the sample characterizing its anisotropy. b) Activating threshold reduction factor for the first 90 neurons with the longest axonal arbor (sorted) for three polarizations of the primary electric field: medial-lateral (red), dorsal-ventral (green), and anterior-posterior (blue). c) Absolute activating thresholds for the same 90 neurons in homogeneous space. d) Absolute activating thresholds for the same 90 neurons within the sample. The units in c,d) are given in terms of the base field strength of 100 V/m; one unit corresponds to 100 V/m. These results are given for the case of the unmyelinated axonal arbor and a negative biphasic activation pulse with the duration of 0.3 ms (Fig. S3).

**Fig. S6.**
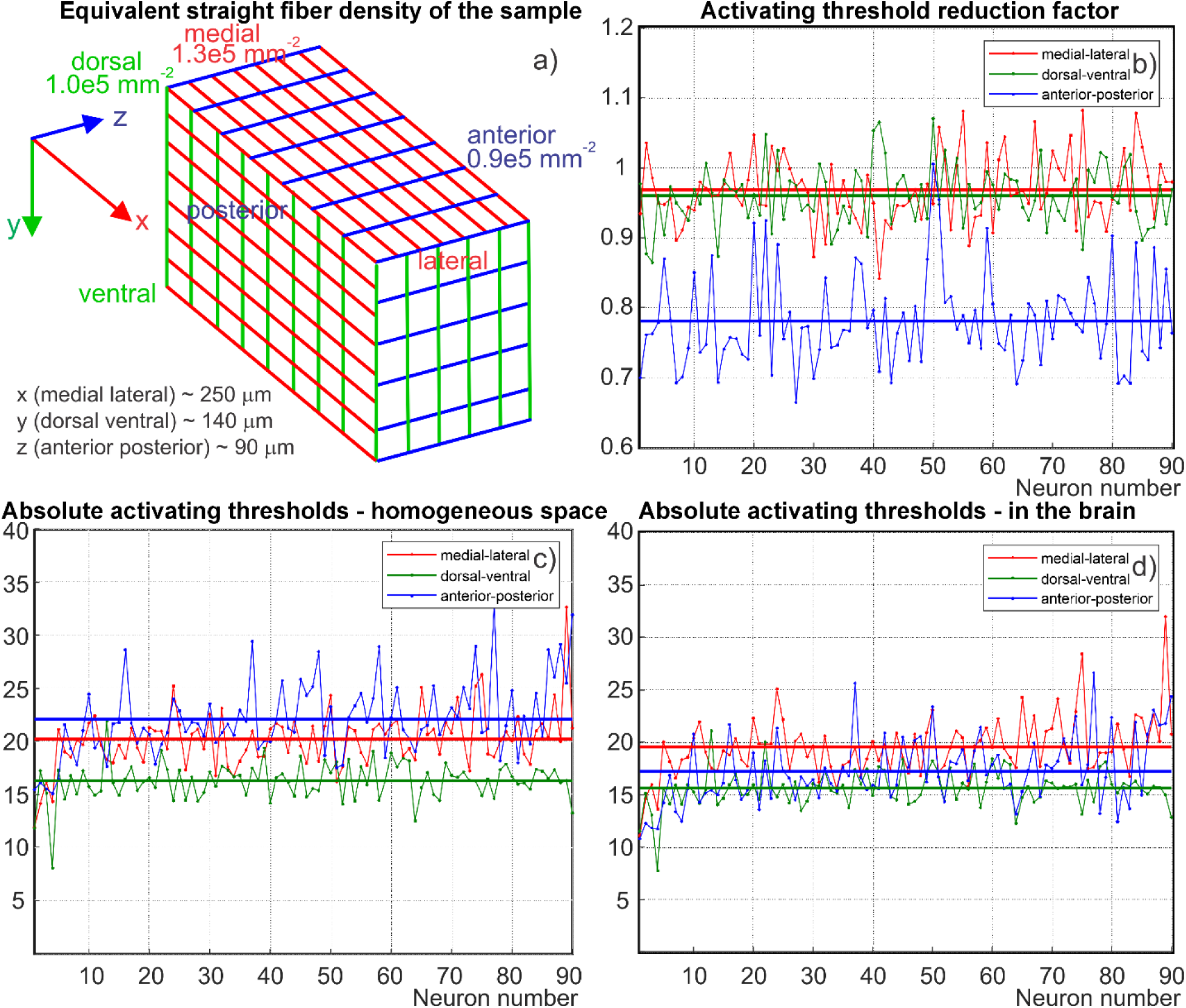
a) Equivalent straight fiber density distribution of the sample characterizing its anisotropy. b) Activating threshold reduction factor for the first 90 neurons with the longest axonal arbor (sorted) for three polarizations of the primary electric field: medial-lateral (red), dorsal-ventral (green), and anterior-posterior (blue). c) Absolute activating thresholds for the same 90 neurons in homogeneous space. d) Absolute activating thresholds for the same 90 neurons within the sample. The units in c,d) are given in terms of the base field strength of 100 V/m; one unit corresponds to 100 V/m. These results are given for the case of the unmyelinated axonal arbor and a positive biphasic activation pulse with the duration of 0.3 ms (Fig. S3).

**Fig. S7.**
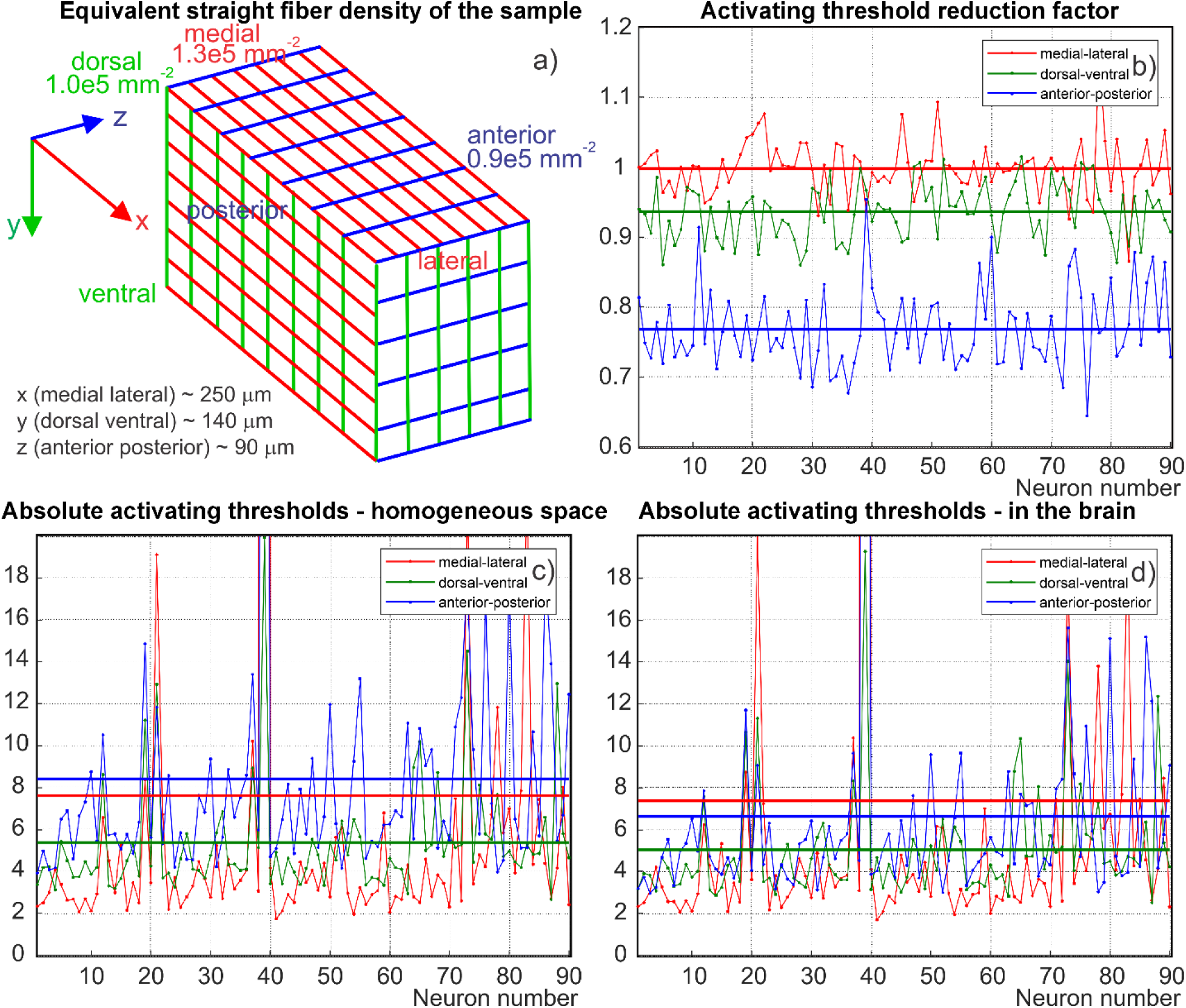
a) Equivalent straight fiber density distribution of the sample characterizing its anisotropy. b) Activating threshold reduction factor for the first 90 neurons with the longest axonal arbor (sorted) for three polarizations of the primary electric field: medial-lateral (red), dorsal-ventral (green), and anterior-posterior (blue). c) Absolute activating thresholds for the same 90 neurons in homogeneous space. d) Absolute activating thresholds for the same 90 neurons within the sample. The units in c,d) are given in terms of the base field strength of 100 V/m; one unit corresponds to 100 V/m. These results are given for the case of the myelinated axonal arbor and a negative rectangular activation monophasic pulse with the duration of 0.1 ms.

**Fig. S8.**
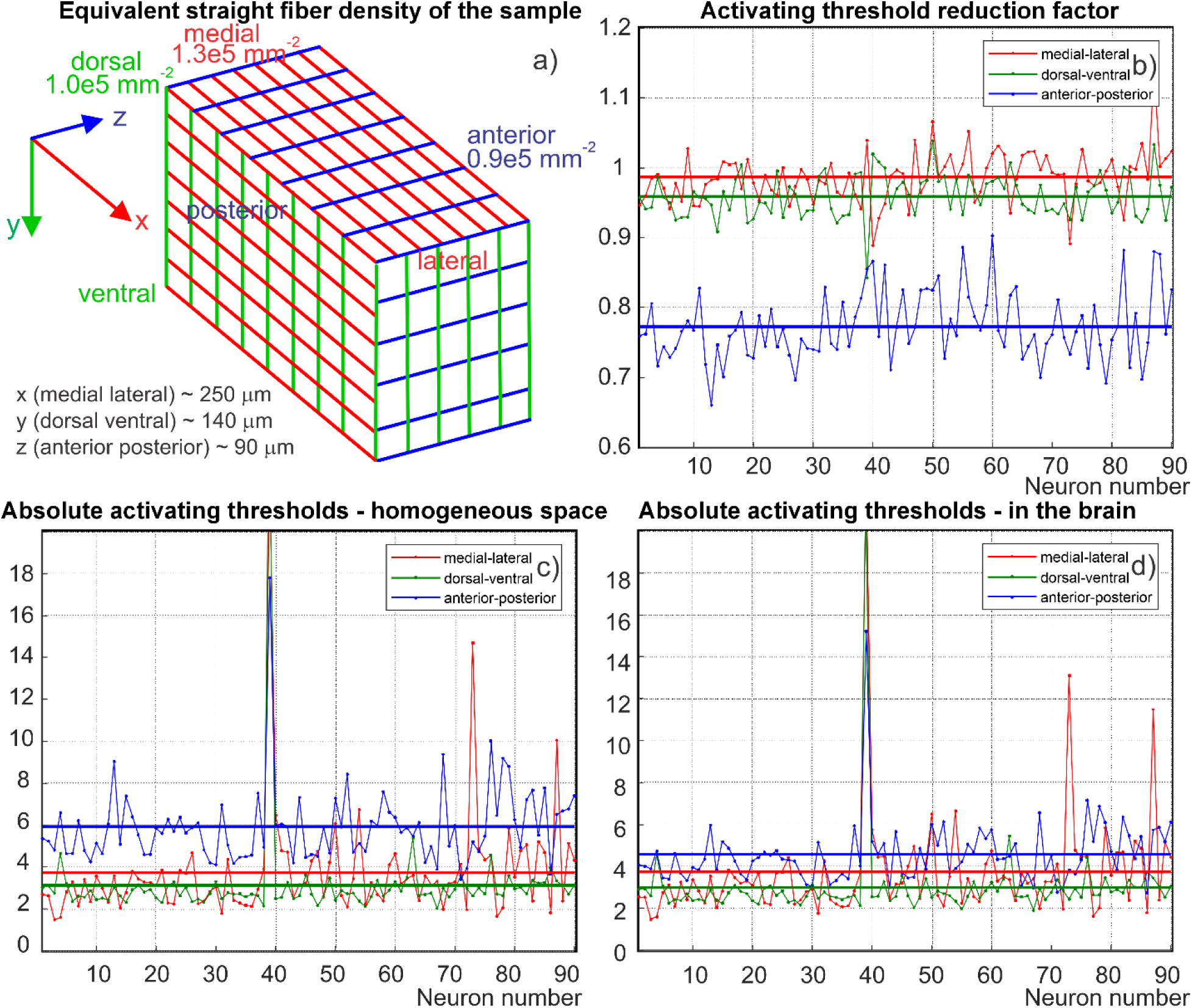
a) Equivalent straight fiber density distribution of the sample characterizing its anisotropy. b) Activating threshold reduction factor for the first 90 neurons with the longest axonal arbor (sorted) for three polarizations of the primary electric field: medial-lateral (red), dorsal-ventral (green), and anterior-posterior (blue). c) Absolute activating thresholds for the same 90 neurons in homogeneous space. d) Absolute activating thresholds for the same 90 neurons within the sample. The units in c,d) are given in terms of the base field strength of 100 V/m; one unit corresponds to 100 V/m. These results are given for the case of the myelinated axonal arbor and a positive rectangular activation monophasic pulse with the duration of 0.1 ms.

**Fig. S9.**
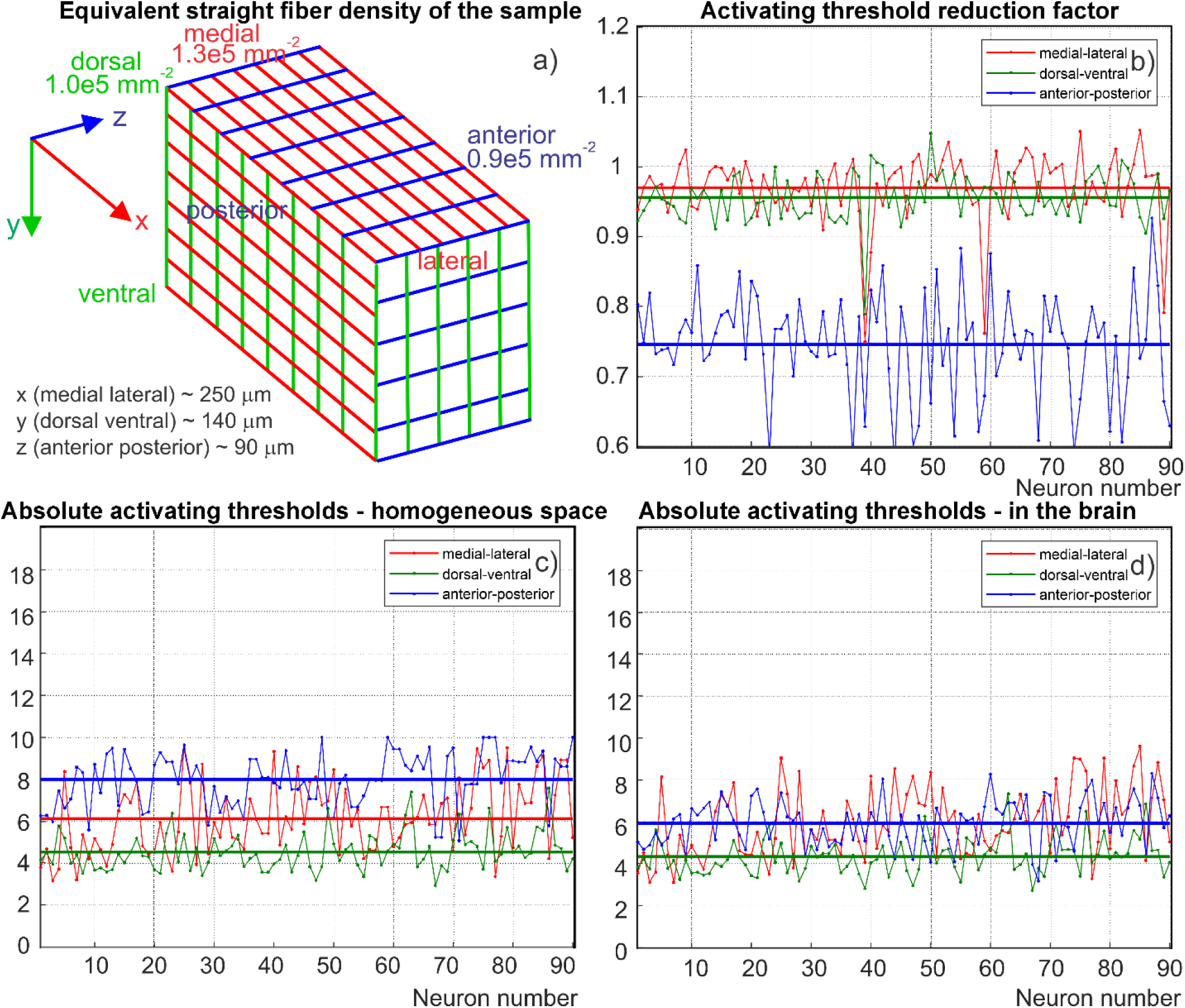
a) Equivalent straight fiber density distribution of the sample characterizing its anisotropy. b) Activating threshold reduction factor for the first 90 neurons with the longest axonal arbor (sorted) for three polarizations of the primary electric field: medial-lateral (red), dorsal-ventral (green), and anterior-posterior (blue). c) Absolute activating thresholds for the same 90 neurons in homogeneous space. d) Absolute activating thresholds for the same 90 neurons within the sample. The units in c,d) are given in terms of the base field strength of 100 V/m; one unit corresponds to 100 V/m. These results are given for the case of the myelinated axonal arbor and a negative biphasic activation pulse with the duration of 0.3 ms (Fig. S3).

**Fig. S10.**
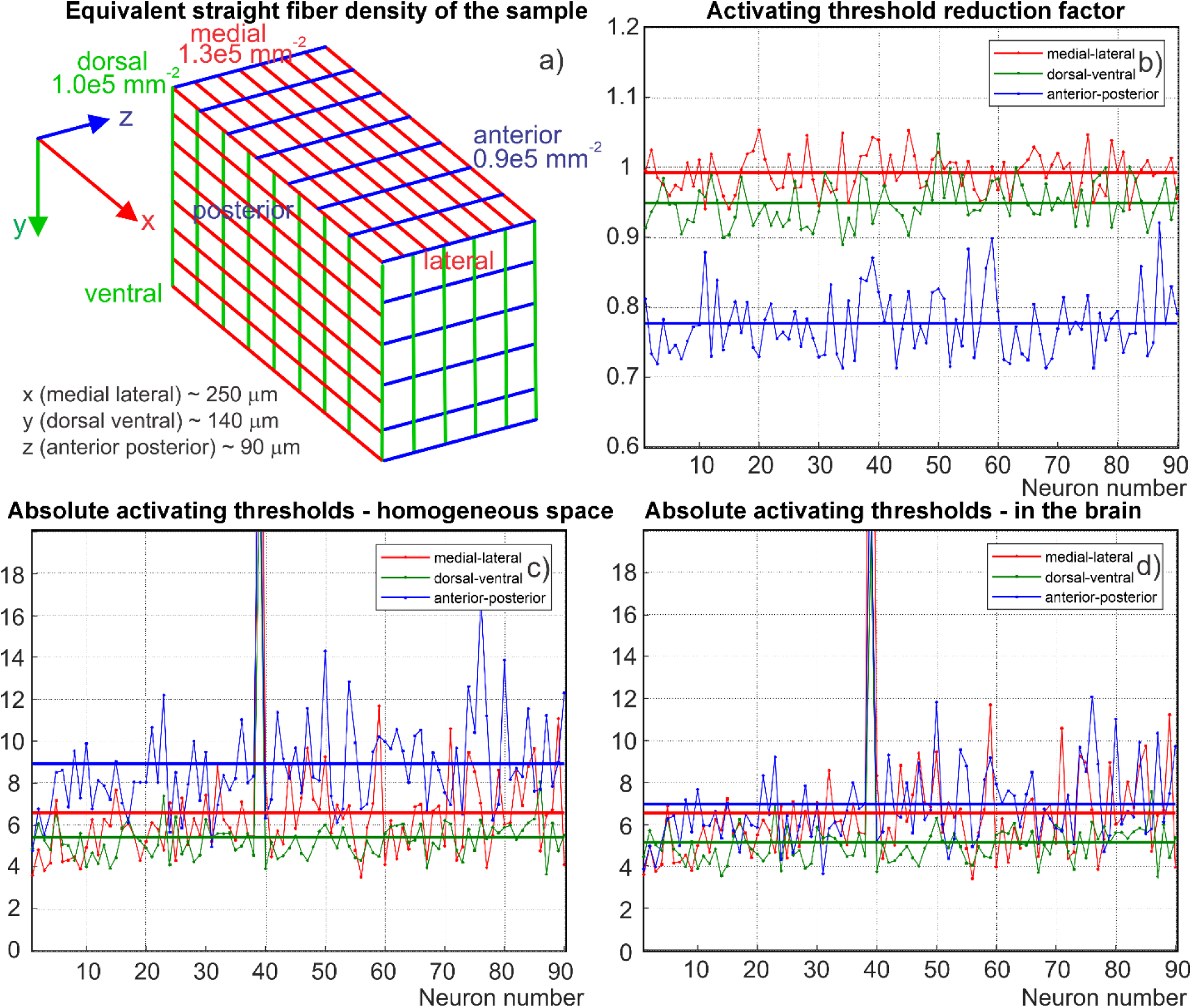
a) Equivalent straight fiber density distribution of the sample characterizing its anisotropy. b) Activating threshold reduction factor for the first 90 neurons with the longest axonal arbor (sorted) for three polarizations of the primary electric field: medial-lateral (red), dorsal-ventral (green), and anterior-posterior (blue). c) Absolute activating thresholds for the same 90 neurons in homogeneous space. d) Absolute activating thresholds for the same 90 neurons within the sample. The units in c,d) are given in terms of the base field strength of 100 V/m; one unit corresponds to 100 V/m. These results are given for the case of the myelinated axonal arbor and a positive biphasic activation pulse with the duration of 0.3 ms (Fig. S3).

## Supplement D Estimates of absolute TMS activating threshold of mouse primary motor cortex from experimental data

### D1. Experimental procedure

In vivo, TMS experiments were performed on male C57BL/6J mice (n¼6) using a custom rodent TMS coil with a magnetic core (Fig. S11c). Animals were anesthetized with sodium pentobarbital intraperitoneally (50 mg/kg), MEP (motor evoked potential) was recorded using a similar approach reported by Rotenberg et al. [13]. All procedures were approved by National Institute on Drug Abuse/NIH (NIDA) animal care and use committee. The coil was mounted to a customized three-axis micromanipulator.

The focal electrical field point was carefully aligned to the targeted mouse motor cortex. Fine adjustments of the coil were made to induce limb twitch on the contralateral hindlimb, but not the ipsilateral hindlimb, any of the forelimbs or any other body part. More detailed description of TMS experiments is given in [14]. Detailed description of a similar custom focal TMS rodent coil was previously given in Ref. [15].

### D2. Modeling setup

Detailed brain model [16] (Fig. S11a) along with the model of the custom TMS rodent coil with a magnetic core (Fig. S11c) has been used to simulate the electric field distribution both in three principal planes (Fig. S11b) as well as just inside the brain compartments (Fig S11d). The skin shell conductivity was chosen as 0.3 S/m, the skull conductivity – 0.01 S/m. All brain compartments were assigned the conductivity of 0.3 S/m. Some conductivity variations were studied. They did not substantially alter the computational results. In general, the small mouse body quite significantly (by a factor of ∼2 or so) blocks the primary applied solenoidal electric field of the TMS coil.

### D3. Estimating electric field strength within mouse brain from measured motor evoked potentials

Simulations were performed for two experimental parameters – the coil current of ∼3,000 A and dI/dt of 94 A/µs – corresponding to the induced MEPs. Fig. S11d shows the resulting magnitude of the total electric field just inside the brain compartments. The circle approximately labels the area of the primary motor cortex. One observes that the field strength there approaches 90 V/m. Conductivity variations of the head compartments lead to the field strength changes within the M1 area (white circle in Fig. S11) to within ±15% about this value. Similar results have been obtained for the total field magnitude distributions in three principal planes shown in Fig. S11e. Another mouse CAD model [17] has also been tested; it also generated very similar results.

**Fig. S11.**
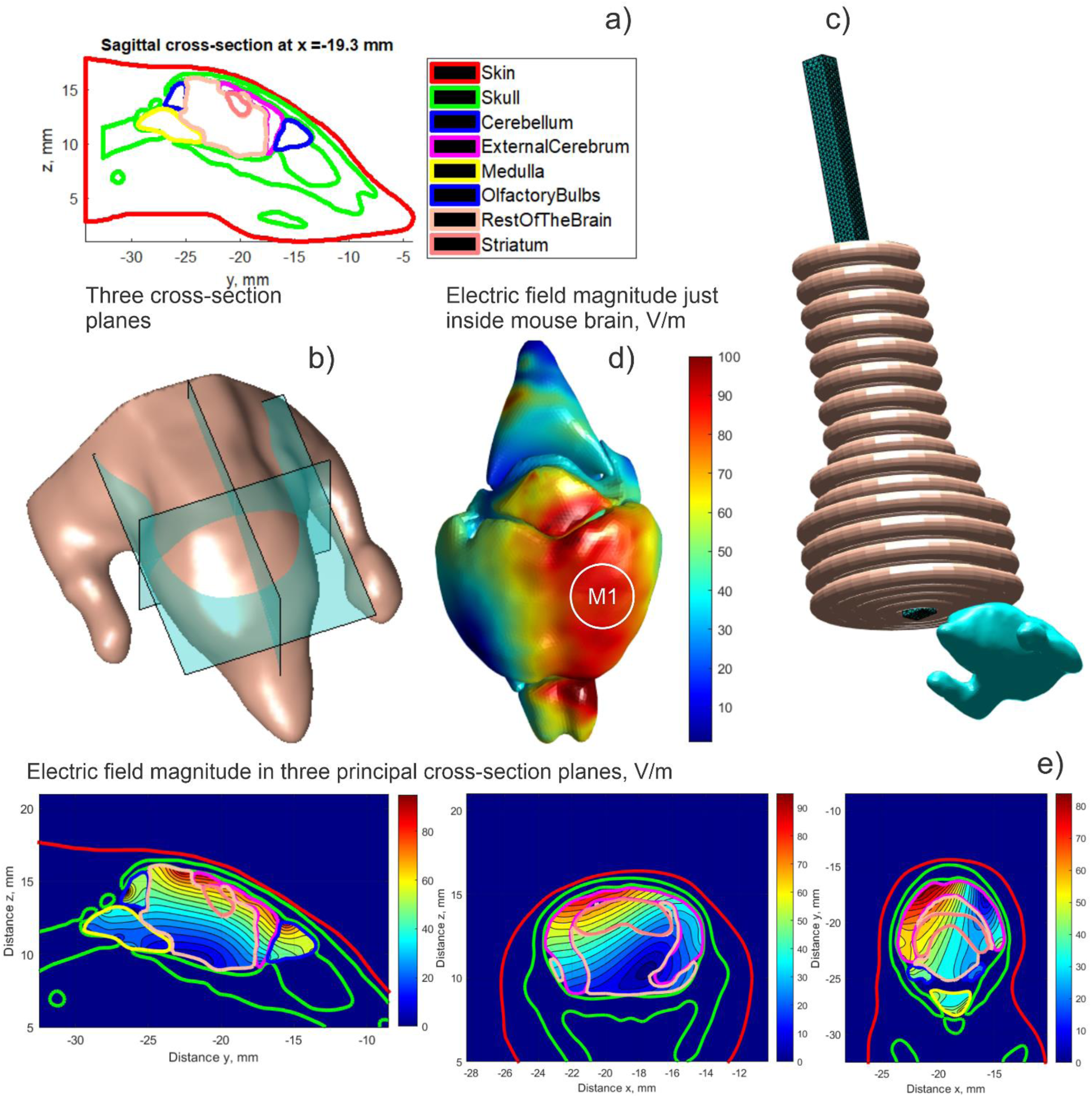
a) CAD Mouse model [16]. b) Three principal planes for field output. c) Shape and positioning the custom TMS coil. d) Electric field magnitude distribution just inside mouse brain. Circle approximately indicates the primary motor cortex.

### D4. Comparison with previously reported computed data for mouse primary motor cortex

In Ref. [17], the maximum fields strengths of 81.3 V/m,113.4 V/m, and 86.5 V/m in the brain have been computed for three different coil positions of a 25 mm figure-8 coil for the coil’s dI/dt of 100 A/μs In Ref. [18], it was concluded from numerical experiments that, for 100 A/μs, the predicted fields in mouse brain are on the order of 120 V/m.

Our experimental data agree with these modeling predictions. They suggest that 90 V/m ± 15% is a reasonable MEP estimate in a mouse brain.

## Supplement E Verification of nested iterative algorithm

### E1. Test sample under study

For comparison purposes, ten nearest neighbors of neuron #192 have been selected as shown in Fig. S12 below. The entire membrane mesh of the sample includes 16 million facets. After that, two methods have been applied:

i. Global iterative BEM-FMM algorithm with the relative residual of 1e-4;
ii. Nested iterative BEM-FMM algorithm of the present study with the total number of 6 neuron-to-neuron outer iterations (as shown in Fig. 3 of the main text) and with the relative residual of 1e-4 for the inner iterative solution.

The y-polarized primary electric field of 100 V/m (corresponding to a vertical direction in Fig. S12) has been applied in both cases. The global algorithm converges after 15 iterations, in ∼1,000 seconds, when using a 3 GHz workstation.

**Fig. S12.**
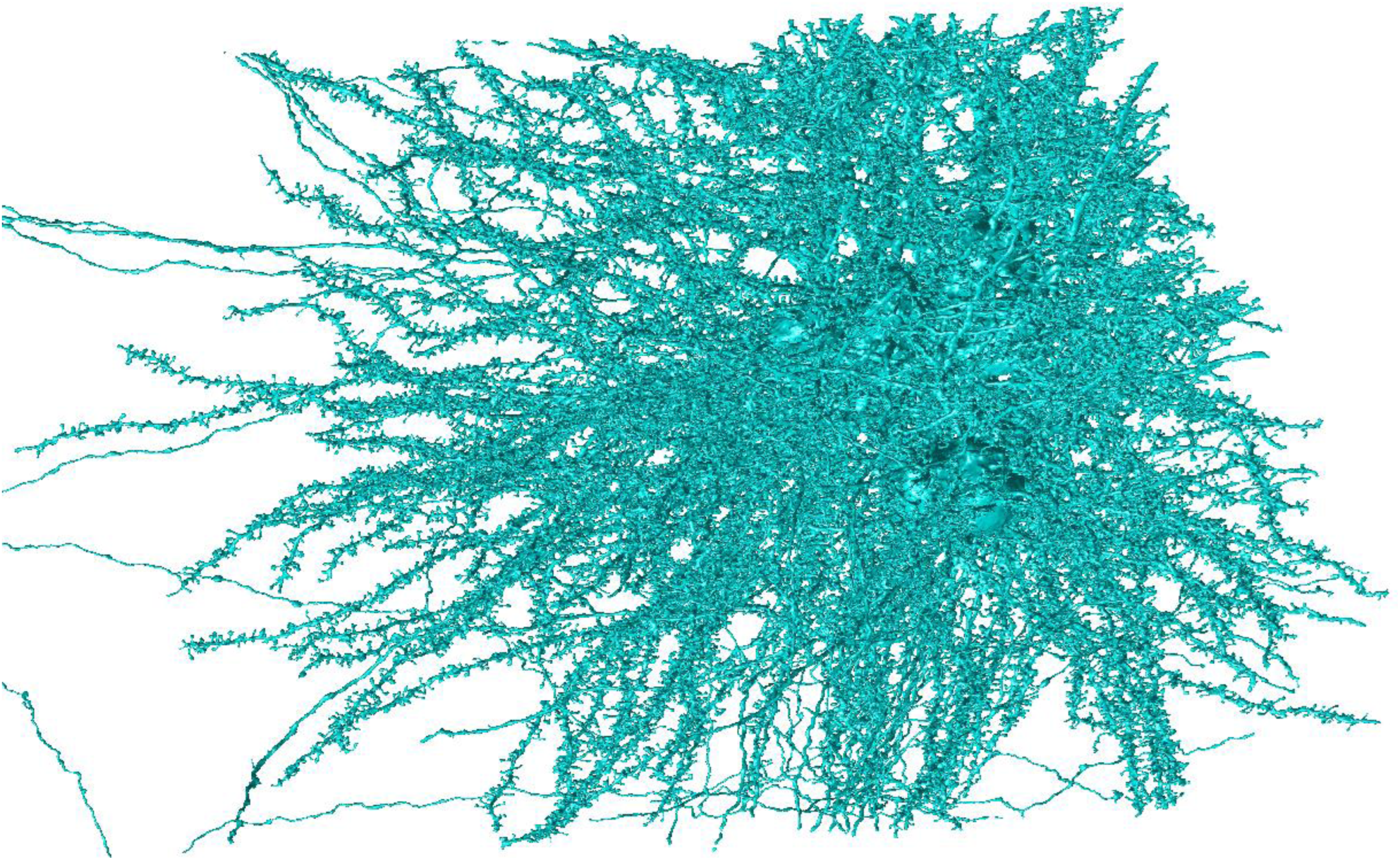
Truncated sample mesh of 11 neighbors of neuron #192.

### E2. Comparison results

The two solutions have been compared to each other with regard to the induced membrane charge densities. Fig. S13 shows the relative 2-norm error in the local charge density for every neuron (11 in total) separately. One observes that this error does not exceed 1% on average.

**Fig. S13.**
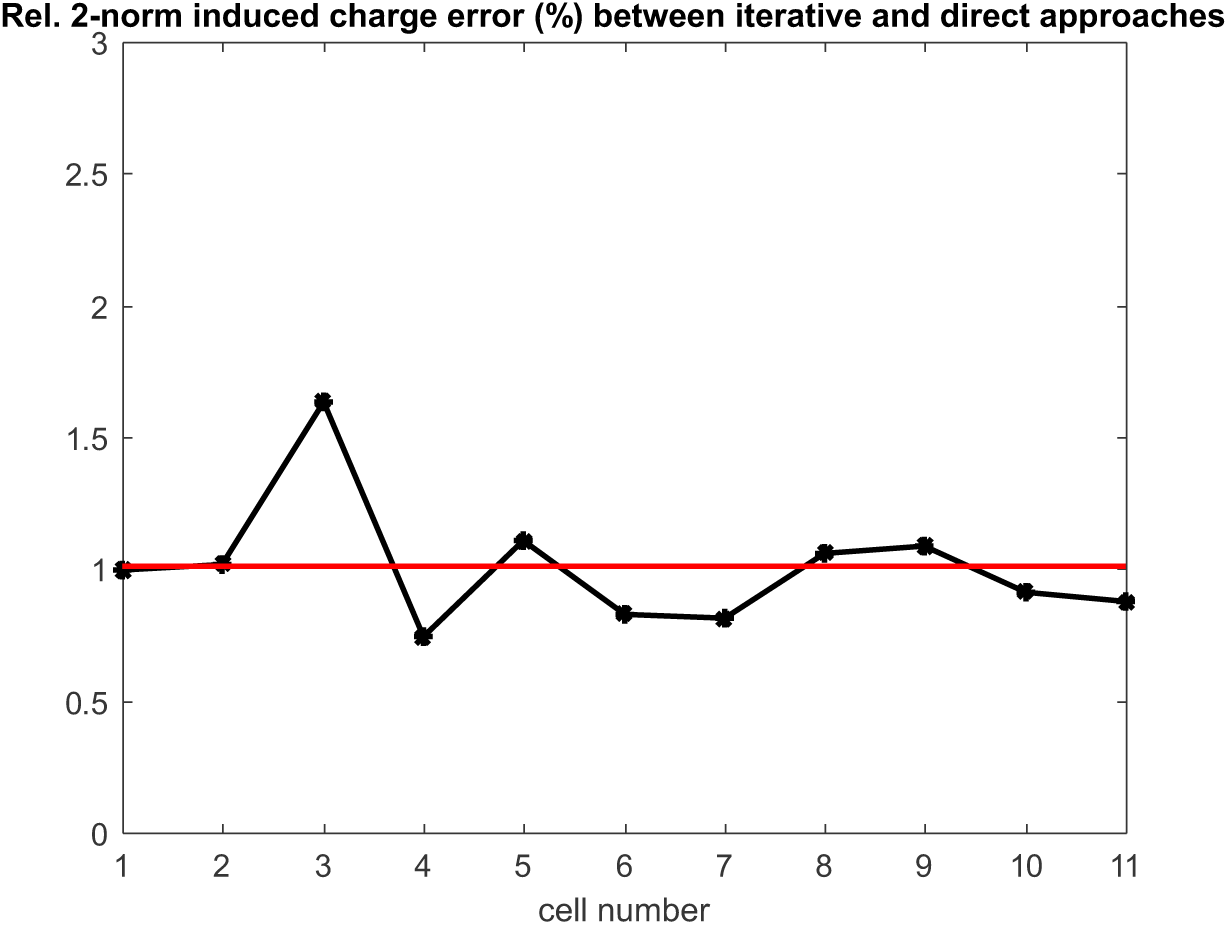
Relative 2-norm error for the induced charge density between the two solutions for every individual neuron of the sample.

## Supplement F Computer code for cathodic stimulation

### F1. Computer code for cathodic stimulation + noise with arbitrarily varying parameters

This code supports Section 4.3 (Discussion Section) of the main text. It is written within MATLAB platform and should run on Windows machines.

A simple numerical experiment with a Hodgkin-Huxley axon originally studied by Rattay [19] is considered. A cathodic stimulation of the 10 mm long axon by a point electrode located at the origin is modeled. The default axonal radius is 5 µm, the length constant λ is 1.5 mm; the time constant τ is 3.3 ms, the resting potential is – 65 mV. A monophasic electric pulse (T=0.1 ms) is applied. Field perturbers are 2D cylinders with non-conducting walls located perpendicular to the axon and with a large radius of 200 µm, chosen for better visualization. An analytical solution for every infinite 2D cylinder secondary field is employed (cf., for example, [20]).

First, the MATLAB script “script01_define_axon_HH_model.m” should be executed. There, the major biophysical and computational parameters are defined. All units are SI units. Extensive comments are provided for every quantity of interest.

Second, the MATLAB script “script02_obtain_axon_solution.m” should be executed. This script runs biophysical computations for the transmembrane potential and the ionic transmembrane current density. If necessary, the total transmembrane current density could be plotted as well.

Two biophysical solutions are executed simultaneously: the first one is for the homogeneous medium (red color) and the second one is for the perturbed extracellular field (blue color). Fig. 9 of the main text shows the graphical output of the MATLAB script “script02_obtain_axon_solution.m” that is updated at every time moment.

Solution parameters (electrode current, axon data, time course, spatial and temporal resolutions, cylinder radii, their spacing and positions) as well as the output format can be modified using both executable MATLAB scripts. Other MATLAB functions should not be changed.

### F2. Code availability

The Dropbox link [21] contains a fully executable computational module built within the MATLAB platform under Windows and a short guide that replicates the above example. This enables both testing of different polarization directions as well as extraction of collinear fields along any desired axonal path.

